# Reductive dechlorination of 2,3-dichloroaniline by *Dehalobacter* in an anaerobic enrichment culture

**DOI:** 10.1101/2025.02.21.639126

**Authors:** Shuping Wang, Sofia P. Araújo, Line Lomheim, E. Erin Mack, Elizabeth A. Edwards, Elodie Passeport

## Abstract

1

2,3-dichloroaniline (2,3-DCA) has widespread use in chemical manufacturing and remains a persistent groundwater contaminant. To better understand the pathway and kinetics of its reductive dechlorination, we conducted a laboratory kinetic experiment using an anaerobic enrichment culture dominated by *Dehalobacter*. At an initial field-relevant concentration of 40 mg/L, complete and stoichiometric dechlorination of 2,3-DCA to aniline via 2-CA was achieved. The intermediates, 2-chloroaniline (2-CA) and 3-chloroaniline, were transiently formed in a ratio of 8:1. The growth yields of *Dehalobacter* on 2,3-DCA and 2-CA were 1.2 ± 0.1 × 10^8^ and 1.3 ± 0.1 × 10^8^ 16S rRNA gene copies/μmol chloride, respectively. The maximum specific growth rate for 2,3-DCA, μ_max_, was 0.18 ± 0.03 day⁻¹ with a half-saturation constant, K_s_, at 45 ± 16 mg /L. The first-order decay constant for *Dehalobacter* when starved of chlorinated electron acceptors was estimated at 0.017 ± 0.001 day⁻¹. Lactate fermentation, acetogenesis from ethanol, syntrophic propionate oxidation, and hydrogenotrophic methanogenesis were observed during dechlorination. This work provides insights into the organohalide respiration of 2,3-DCA to aniline and advances the understanding of microbial interactions during anaerobic dechlorination. These results offer guidance for developing stable dechlorinating microbial ecosystems and key kinetic parameters for predictive modeling of groundwater 2,3-DCA fate and transport.

**Synopsis:** This study demonstrated stoichiometric dechlorination of 2,3-dichloroaniline to aniline, characterized syntrophic interactions in a mixed anaerobic dechlorinating culture, and determined growth yield, first-order decay and Monod constants for *Dehalobacter*.

**Graphical abstract:** 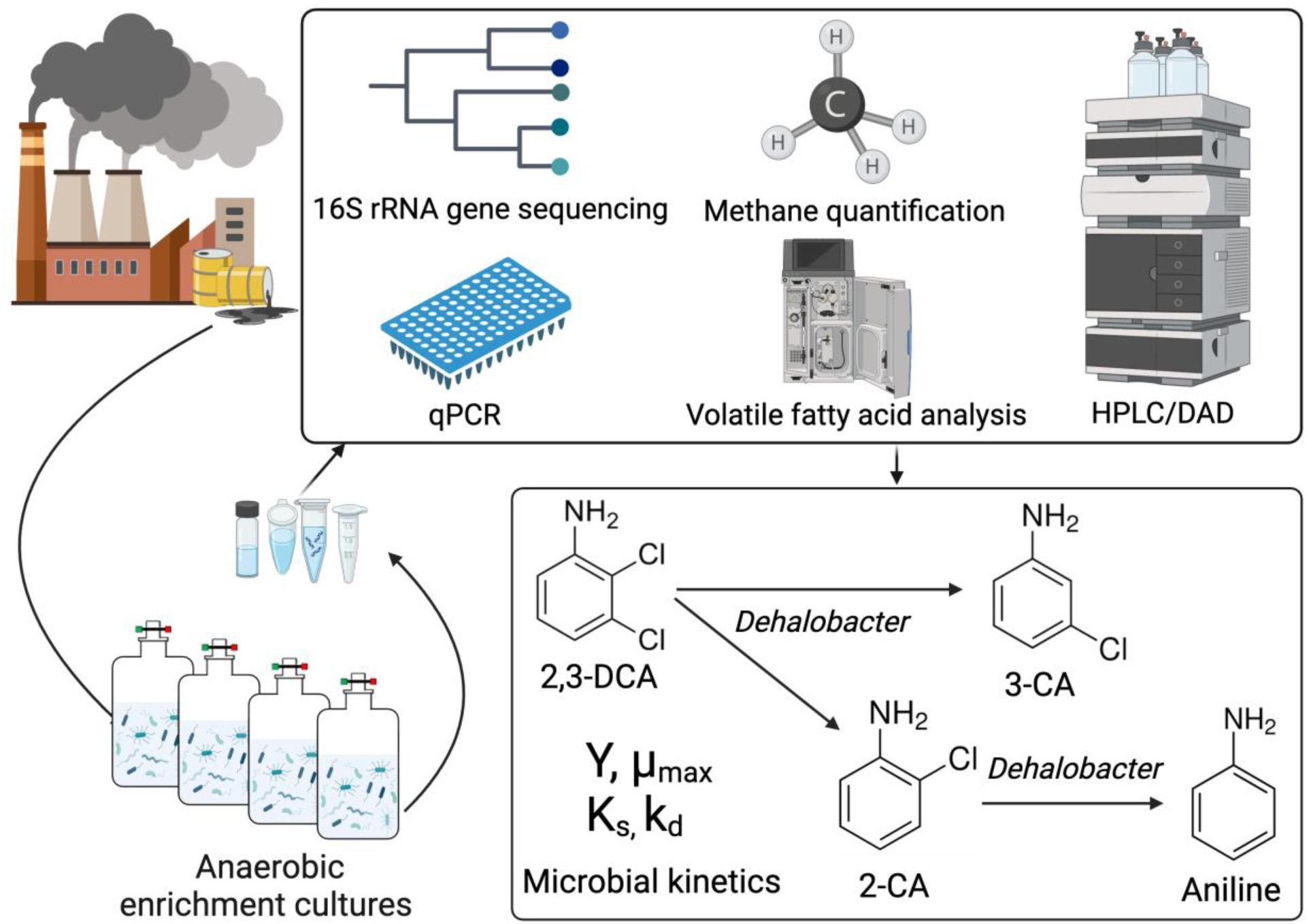

Created in BioRender. Wang, S. (2025) https://BioRender.com/f47t993

## 4 INTRODUCTION

Dichloroaniline (DCA) isomers are a class of amino-substituted chlorinated aromatic compounds that are widely used as intermediates in the manufacture of various chemical products, including herbicides, cosmetics, pigments, pharmaceuticals, and antimicrobials.^1^ DCAs are primarily released into the environment through improper handling and disposal during industrial processes. They are also produced from natural attenuation of phenylurea herbicides and nitro-substituted aromatics.^2–4^ Due to high chemical stability and toxicity, DCA contamination poses a significant environmental challenge, particularly in aquatic and soil ecosystems.^5–7^

Contaminated groundwater environments, which are often anoxic, provide favorable conditions for organohalide respiration. Such processes have proven successful for many halogenated pollutants, particularly chlorinated alkenes and alkanes.^8–12^ However, studies on the reductive dechlorination of DCAs remain limited.

Reductive dechlorination of DCAs was first observed using pond sediment^13^ and aquifer slurries^14^. The latter study reported the biotransformation of 3,4-DCA (around 50 μM) to 3-chloroaniline (3-CA) during the 8-month experiment.^14^ Similarly, Susarla et al. (1998) observed reductive dechlorination of 3,4-DCA (4 μM) using sulfidogenic estuarine sediment with 3-CA and aniline as the end products.^15^ However, these early studies used low DCA concentrations and often required extended incubation periods (e.g., >200 days) for complete transformation.

Moreover, the microorganisms responsible for these anaerobic transformations were not identified and the relatively poor mass balance obtained in these studies limited the understanding of the underlying organohalide respiration pathways.

More recent studies have identified microorganisms capable of organohalide respiration using DCAs. Duc and Oanh (2019) isolated *Geobacter* sp. KT5 from contaminated river sediment, which grew on a wide range of chlorinated anilines (50 μM) and transiently produced dechlorinated metabolites such as 3-CA, aniline and 4-aminobenzoate.^16^ Cooper et al. (2015) showed that organohalide respiring *Dehalococcoides* strain CBDB1 could respire and dechlorinate 2,3-DCA (but not 2,4- or 2,6-DCA) with H_2_ as electron donor.^17^ In a follow up study contrasting dechlorinating mechanisms between *Dehalococcoides mccartyi* strain CBDB1 and *Dehalobacter* sp. strain 14DCB1 previously isolated on chlorinated benzenes, the authors found that both strains could transform 2,3-DCA and 3,4-DCA with hydrogen as the electron donor producing 3-CA (by CBDB1) or very minor amounts of 2-chloroaniline (2-CA, by 14DCB1) and 3-CA (by 14DCB1).^18^ Despite these advances, no studies have focused on achieving complete and stoichiometric anaerobic transformation of DCAs to aniline at concentrations relevant to those observed at a highly contaminated site^19–21^ with average total DCA concentrations above 200 μM in groundwater. Furthermore, although *Dehalobacter* can transform a variety of halogenated organic contaminants (e.g., chlorinated ethanes, toluene, benzenes, and chloroform),^22–24^ reproducible and robust microbial kinetics of *Dehalobacter*-mediated dechlorination are lacking, limiting the predictability and quantification of natural or bioaugmented dechlorination at contaminated sites. The interactions between *Dehalobacter* and other microorganisms in dechlorinating microbial consortia also remain poorly understood. In a companion study, subtle aniline formation was observed after 2,3-DCA dechlorination to 2-CA in culture 23DCA-T2.^25^ We made a transfer from this culture and the resulting subculture surprisingly exhibited significantly more aniline formation. This prompted us to further explore dechlorination of 2,3-DCA to aniline and *Dehalobacter* kinetics. The objectives of our study were to: (1) demonstrate the complete reductive dechlorination of 2,3-DCA to aniline via 2-CA at a high initial concentration; (2) identify the associated transformation pathway; (3) determine growth yield and kinetic parameters to improve the predictability of dechlorination and (4) investigate syntrophic interactions between cocultured microorganisms and *Dehalobacter*. These objectives were achieved by frequent (every 3 days) sampling of chemical and microbial concentrations in stable enrichment cultures derived from a contaminated site.

## 5 MATERIALS AND METHODS

### 5.1 Chemicals

All chemicals were obtained from Sigma Aldrich (St. Louis, Missouri, USA) unless otherwise noted. Cultures were amended with 2,3-DCA (99.6% purity). Aniline and 2-CA used as standards were both at 99.5% purity; whereas, 3-CA and 4-chloroaniline (4-CA) were 99% and 98% pure, respectively. Ethanol was HPLC grade. Sodium lactate (60% by mass) was purchased from Spectrum Chemical MFG Corp.

### 5.2 Microbial Enrichment Cultures

The culture used as an inoculum in this study was the third consecutive subculture (referred to as WANG-23DCA-T3) from an anaerobic methanogenic 2,3-DCA-dechlorinating enrichment culture (i.e., 23DCA-T2), described in Araújo et al. (2025).^25^ In brief, 23DCA-T2 originated from microcosms established in 2015 using groundwater and soil samples collected from a heavily contaminated industrial site in Brazil (Supporting Information (SI) Figure S1).^25^ On May 4, 2022, the first transfer (WANG-23DCA-T1) was prepared by inoculating 90 mL of anaerobic medium with 10 mL 23DCA-T2 sample. After three-month cultivation, on August 9, 2022, a 50 mL portion of culture WANG-23DCA-T1 was transferred to establish a new subculture (WANG-23DCA-T2). On day 188 of WANG-23DCA-T2 (February 13, 2023), a 3% transfer of culture (12 mL) from WANG-23DCA-T2 was made to establish subculture WANG-23-DCA-T3 (400 mL). All three subcultures were repeatedly amended with 2,3-DCA and demonstrated activity in transforming it, with WANG-23-DCA-T3 showing particularly strong conversion of 2,3-DCA to aniline. These subcultures were topped up with medium multiple times to build up culture volume for future experiments. On day 254 of WANG-23-DCA-T3 (October 25, 2023), the inoculum for this study was taken from WANG-23-DCA-T3. Dechlorination profiles and associated cultivation conditions for subcultures and transfers are detailed in SI Section S1 and Figures S2-S4.

### 5.3 Experimental Setup

In this study, all experiments were performed in an anaerobic glovebox (Vinyl Anaerobic Chamber glovebox, Coy Laboratory Products) at room temperature (21-22°C). The glovebox was filled with a gas mix of 10% H_2_, 10% CO_2_ and 80% N_2_, however owing to reaction with diffusing oxygen, the H_2_ concentration in the glovebox remained around 1-1.5% H_2_.

To establish the dechlorination kinetics, four replicates with an initial concentration of 40 mg/L (i.e., 250 μM) 2,3-DCA and 3% (v/v) inoculum were prepared in autoclaved 250-mL Boston round glass bottles sealed with Mininert valves, with an initial medium volume of 210 mL and headspace volume of 40 mL. They are referred to as “biotic bottles”. The medium used in this study was a pre-reduced mineral medium described in Edwards and Grbic-Galić (1992).^26^ To estimate the decay kinetics of *Dehalobacter* under conditions lacking electron acceptors, triplicate decay bottles were established in the same manner as the biotic bottles, excluding the addition of 2,3-DCA. To monitor the mass loss due to sampling and control for the possibility of abiotic transformation processes, triplicate abiotic controls were set up similarly without inoculation in autoclaved 250-mL Boston round bottles, each containing 210 mL of mineral medium spiked at 40 mg/L of 2,3-DCA. Biotic and abiotic bottles were amended with 2,3-DCA by dropping a feeding vial containing 2,3-DCA into the bottle. Details are provided in Section S2. After spiking with 2,3-DCA, the concentration of 2,3-DCA was regularly monitored in all bottles. Inoculation of the biotic bottles was not conducted until the concentration of 2,3-DCA had stabilized at the target concentration (i.e., 40 mg/L), which took three days. Once complete dissolution of 2,3-DCA was attained, 6.5 mL of homogenized, active inoculum was added to four biotic bottles. The biotic and decay bottles were then amended with sodium lactate (0.6 mM) and ethanol (3.3 mM). Abiotic controls did not receive any amendment of electron donors. The inoculation defined the time zero of the experiment. Samples were collected immediately after inoculation to characterize the initial concentration of methane (headspace), 2,3-DCA, transformation products, volatile fatty acids, inorganic anions, *Dehalobacter* abundance, and microbial community composition. During the experiment, all the bottles were covered with black cloth to minimize exposure to light and possible photodegradation.

Aqueous samples (350 µL) were taken from biotic and abiotic bottles approximately three times every week for chloroaniline and aniline analysis. A 4-mL sample was collected for DNA extraction from biotic and decay bottles roughly every ten days. Samples (500 µL) for quantifying inorganic anions and volatile fatty acids were taken weekly from biotic and abiotic bottles. Headspace samples (300 µL) for determining methane production were withdrawn approximately once every ten days from the biotic bottles. The total liquid volume withdrawn did not exceed 20% of the initial volume during the first 50 days (i.e., dechlorination of 2,3-DCA to 2-CA and 3-CA). Sample collection and processing are detailed in the SI.

### 5.4 Analytical procedures

Chloroanilines and aniline were measured using high-performance liquid chromatography coupled with a diode array detector (HPLC/DAD, Thermo Scientific Dionex Ultimate 3000 series) equipped with an Accucore C18 column (100 mm × 2.1 mm × 2.6 μm) and guard column (10 mm × 2.1 mm × 2.6 μm; Thermo Scientific, Waltham, MA, USA). Methane and other possible volatiles (e.g., benzene) were measured by a Hewlett-Packard 5890 Series II gas chromatograph (GC) using a GSQ I. D. PLOT column (30 m × 0.53 mm; J & W Scientific, Folsom, USA) configured with a flame ionization detector (FID). Inorganic anions (sulfate, phosphate, nitrate, nitrite, and chloride) and volatile fatty acids (VFAs; acetate, propionate, butyrate, lactate, formate, and pyruvate) were measured by ion chromatography (IC) using a Dionex™ Integrion™ high pressure IC coupled with an ASRS 500 suppressor and a Dionex™ AS-DV autosampler (Thermo Fisher Scientific). Separation was achieved using a Dionex™ IonPac™ 2 × 250 mm AS11-HC column. Analytical methods are detailed in Sections S3-S5. The limits of quantification (LOQs) were 0.7 µmol/L for 2,3-DCA, 0.8 µmol/L for 2-CA and 3-CA, 1.5 µmol/L for aniline, 0.003 mmol/L for lactate, and 0.001 mmol/L for all other target VFAs (Section S5).

### 5.5 DNA sampling and extraction

Culture samples (4 mL) were withdrawn for DNA extraction and centrifuged (described in Section S6), supernatant discarded, and the cell pellets were stored at −80 °C. DNA was extracted using the DNeasy PowerSoil Pro kit (Qiagen) following manufacturer’s guidelines. DNA was eluted in 50 μL of 10 mM Tris-HCl buffer and stored at −80 °C until analysis.

### 5.6 Quantitative PCR analysis

Quantitative polymerase chain reaction (qPCR) primer sets targeting *Dehalobacter* (Dhb) were used as previously described.^27^ Primer sequences were Dhb477f (5’-GATTGACGGTACCTAACGAGG-3’) and Dhb647r (5’-TACAGTTTCCAATGCTTTACG-3’).^27^ Primers targeting most bacteria (Bac) were as follows: Bac1055f (5’-ATGGCTGTCGTCAGCT-3’) and Bac1392r (5’-ACGGGCGGTGTGTAC-3’).^28^ Temperature profiles and reaction details are available in Section S7. The qPCR results with *Dehalobacter* biomass estimation are provided in Table S1.

### 5.7 16S rRNA gene amplicon sequencing

16S rRNA gene amplicon sequencing targeting the V6-V8 region of bacteria and archaea was carried out using the Illumina NovaSeq PE300 platform at Genome Québec (Montreal, QC, Canada), using a set of modified “staggered end” primers: forward primer 926f (5’-AAACTYAAAKGAATWGRCGG-3’) and reverse primer 1392r (5’-ACGGGCGGTGWGTRC-3’, as previously described.^29^ Original amplicon reads were processed using QIIME 2 (v.2021.2) (Section S8).^30^ Amplicon sequence variants (ASVs) were generated using DADA2 plugin ^31^ and taxonomically classified with the SSU SILVA v.138 database.^32^ Relative abundance of ASVs in bacterial (Table S2) and archaeal (Table S3) communities were calculated. Sequence alignment for *Dehalobacter* ASVs was performed in Geneious Prime (v.2023.1.2) using the MUSCLE Alignment feature. Original sequence reads are available through NCBI (Accession: PRJNA1040917).

### 5.8 Modeling dechlorination of 2,3-DCA using Monod kinetics

We applied the Monod equations^33,34^ to estimate the kinetic parameters for *Dehalobacter* growth on 2,3-DCA with two coupled non-linear differential equations that describe the growth of *Dehalobacter* (Eq. 1) and the associated consumption of the limiting substrate (Eq. 2):

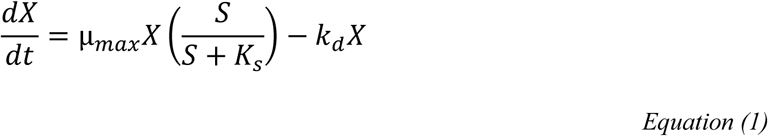

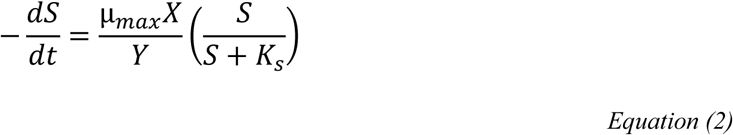

where S and X represent concentrations (g/L) of the substrate (2,3-DCA or 2-CA) and active biomass (*Dehalobacter*) at time t, respectively; Ks is the half-saturation constant (g/L), and µ_max_ is the maximum specific growth rate (day^−1^); Y is the growth yield (g biomass /g substrate); k_d_ is the first-order decay constant (day^−1^) for biomass; dX/dt is the growth rate of biomass (g/L/day), and −dS/dt is the substrate consumption rate (g/L/day).

Yield and decay parameters Y and k_d_ were independently determined in this study. Therefore, K_s_ and µ_max_ were the only two Monod parameters obtained through the least squares optimization. The time-course substrate depletion data were used to fit the Monod model. The coupled differential equations were numerically solved using the “solve_ivp” function from the “scipy.integrate” module in Python, with initial conditions set to the substrate and biomass concentrations measured at time zero (S_0_ and X_0_, respectively). S_0_ was well constrained, but X_0_ being a less accurate measurement was allowed to vary within a narrow range during optimization. The optimal initial biomass for modelling is referred to as X_0_’. The combined residual sum of squares (RSS) from all replicates was minimized. We employed the “least_squares” function from the “scipy.optimize” module in Python to achieve the best fit by adjusting optimization parameters, K_s_, µ_max_, and X ^’^ for 2,3-DCA dechlorination. The starting biomass used for modeling the subsequent 2-CA dechlorination was calculated from the kinetics of the preceding 2,3-DCA dechlorination step. Thus, for 2-CA dechlorination kinetics, only K_s_ and µ_max_ were adjusted to minimize RSS. To accommodate the determination of K_s_ and µ_max_, broad parameter ranges were applied ([0,1000] mg/L for K_s_ and [0,10] day^−1^ for µ_max_). A range of [X_0_/2, 4X_0_] for X_0_’ was set to account for potential estimation errors associated with the initial biomass measurement using qPCR. The goodness-of-fit was assessed using RSS, reduced chi-square, and root-mean-square errors (RMSE).

## 6 RESULTS AND DISCUSSION

### 6.1 Complete dechlorination of 2,3-DCA to aniline via 2-CA

We observed complete and stoichiometric transformation of 2,3-DCA to aniline via 2-CA in all four biotic replicates. The data for Biotic Bottle I is shown in Figure 1a, with the results for the other replicates available in Figure S5. In the abiotic controls, 2,3-DCA concentration remained constant throughout the course of the experiment (Figure 1a) and no transformation products were detected.

**Figure 1.**
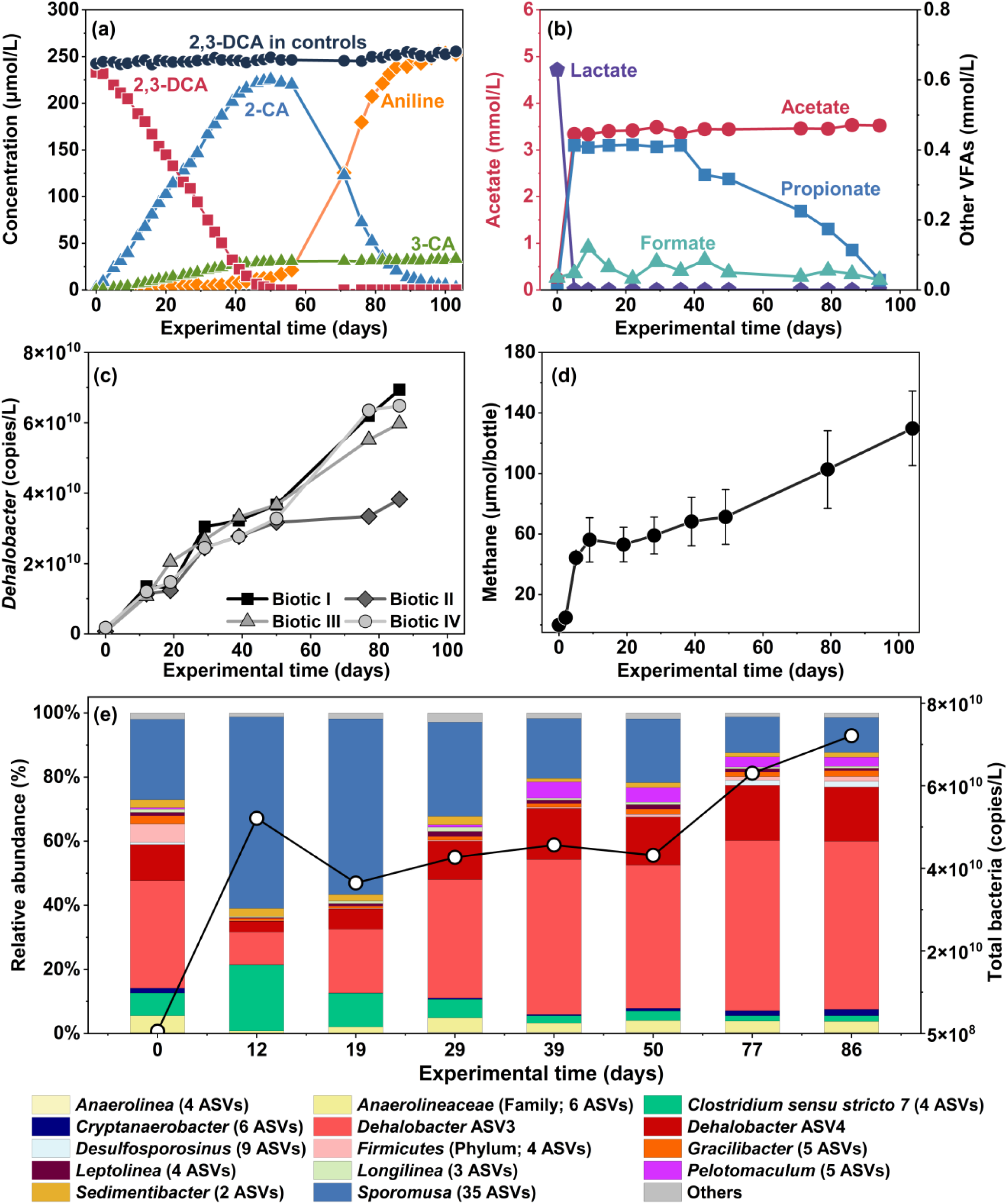
(a) Dechlorination of 2,3-DCA to aniline via 2-CA in biotic replicate I. Concentrations (µmol/L) of 2,3-dichloroaniline (2,3-DCA, red squares), 2-chloroaniline (2-CA, blue triangles), 3-chloroaniline (3-CA, green triangles), and aniline (orange diamonds) over the experimental time (days) (Table S5). 4-CAwas not detected. Aniline and 3-CA are shown as net formation by subtracting their initial concentrations. The average 2,3-DCA concentrations of triplicate abiotic controls are represented by black circles. (b) Concentration (mmol/L) of lactate (purple pentagons), acetate (red circles), propionate (blue squares), and formate (light teal triangles) in replicate I. Butyrate and pyruvate were not detected and fatty acid concentrations in abiotic controls remained below the LOQ (Table S6). (c) Increase in *Dehalobacter* absolute abundance (16S rRNA gene copies/L) in all biotic replicates. The absolute abundance was measured using qPCR with primers targeting *Dehalobacter*. (d) Average methane production (µmol per bottle) of all biotic replicates. Error bars represent the standard deviation among the four biotic replicates (Table S7). (e) Changes in bacterial composition during dechlorination in replicate I. The relative abundance of bacteria at the genus level (unless otherwise specified) obtained from 16S rRNA amplicon sequencing, shown as stacked bars, is presented in combination with total bacterial concentrations (gene copies/L, shown on the secondary y-axis) determined using qPCR targeting general bacteria. Others: ASVs with relative abundance < 1%. Results from all four biotic replicates are provided in Figures S5, S6, S7a, and S7b.

In the biotic bottles, between days 0 and 53, 2,3-DCA underwent quantitative dechlorination without a significant lag phase (< 2 days), resulting in the formation of 2-CA and 3-CA at an overall molar stoichiometric ratio of 7.6 (± 1.9): 1 (n=88 using all time points). Our observations indicate that *Dehalobacter* preferentially removes the chlorine substituent at the meta position relative to the amino group to produce 2-CA. This finding aligns with a previous study showing the dechlorination of 2,3-DCA to 2-CA by *Dehalobacter* Strain 14DCB1.^18^ However, in that study, the formation of 2-CA was minimal (<0.8 µM) over five successive 2,3-DCA feeding cycles at 40 µM across 35 days.^18^ In contrast, we observed a 1:1 stoichiometric conversion of 2,3-DCA to CAs (mainly 2-CA). A mole balance analysis for each biotic replicate showed that 92 (± 4) % (n=4) of the reduction in 2,3-DCA can be explained by the production of dechlorination intermediates 2-CA and 3-CA over the course of 50 days.

After day 50, 2-CA was further dechlorinated to aniline, whereas 3-CA persisted (Figures 1a and S5). End products of this two-step dechlorination were predominantly aniline (89%), with 3-CA constituting the balance. On day 103, complete dechlorination of 2-CA to aniline was observed. Dechlorination of 2-CA was also quantitative as 88 (± 6) % (n=3) of 2-CA consumed can be attributed to the corresponding increase in the aniline concentration. Dechlorination of 2,3-DCA to aniline is a sequential process. Dechlorination of 2-CA to aniline did not start until 2,3-DCA was depleted in the bottles. This observation aligns with the results from the parent culture (Figure S4), which shows that aniline formation began when 2-CA concentration was much higher than 2,3-DCA concentration.

The preferential utilization of 2,3-DCA as an electron acceptor by *Dehalobacter* over 2-CA is consistent with the reported Gibbs free energy under standard conditions for dechlorination of 2,3-DCA to either 2-CA (−135.0 kJ/reaction) or 3-CA (−137.7 kJ/reaction), which are more thermodynamically favorable than the dechlorination of 2-CA to aniline (−126.8 kJ/reaction).^35^ Moreover, this preferential utilization may explain the negligible aniline formation in the earlier subcultures (Figures S2 and S3), where 2,3-DCA was re-amended before its depletion and consistently available to *Dehalobacter*.

### 6.2 *Dehalobacter* is responsible for dechlorination

A continuous and consistent rise in *Dehalobacter* absolute abundance (16S rRNA gene copies/L) was observed across all biotic replicates during both the 2,3-DCA and 2-CA dechlorination stages (Figure 1c). Biotic bottle II showed lower *Dehalobacter* growth during 2-CA dechlorination compared to other replicates, but this aligns with the limited dechlorination of 2-CA to aniline in this bottle (Figure S5II), confirming the correlation between 2-CA dechlorination and *Dehalobacter* growth. The results also indicate that *Dehalobacter* can be enriched through the supply of 2,3-DCA and 2-CA (Figure 1.e). From days 12 to 87, the relative abundance of *Dehalobacter* in the bacterial community increased by 46.9 ± 8.4% (n=4), becoming the most dominant bacterial genus. In line with stoichiometry, the results demonstrate that *Dehalobacter* can grow on 2,3-DCA and 2-CA through reductive dechlorination. Two predominant ASVs (ASV3 and ASV4) accounted for 99.9 ± 0.1% (n=32) of total *Dehalobacter* reads (Table S2). *Dehalobacter* ASV3 and ASV4 shared >99.78% 16S rRNA gene sequence identity (differing by one base pair in the 468-bp amplicon) and exhibited consistent abundance trends throughout the experiment, maintaining a stable ASV3/ASV4 ratio of 3.1 ± 0.1:1 (n=32) in all the samples. Therefore, the two ASVs likely belong to the same *Dehalobacter* strain.

*Dehalobacter* ASV3 and ASV4 were not the dominant *Dehalobacter* ASVs (ASV1 and ASV2) in the original parent culture (23DCA-T2)^25^ but the ASVs were present in other enrichment cultures stemming from the original site material. With approximately 97% sequence identity (Table S4), both ASV3 and ASV4 are genetically distinct from the dominant ASV1 and ASV2 found in 23DCA-T2,^25^ where slow and low-level aniline formation was noticed. While these genetic differences may account for the distinct capability of using 2-CA as an electron acceptor to yield aniline, future research should investigate this further.

### 6.3 Growth yields and microbial kinetics of *Dehalobacter*

#### 6.3.1 Growth yields and first-order decay

Through the two-step dechlorination process, we measured *Dehalobacter* growth yields on 2,3-DCA and 2-CA (Figure 2). The growth yield for 2,3-DCA dechlorination was 1.2 ± 0.1 × 10^8^ 16S rRNA gene copies/μmol 2,3-DCA converted (Figure 2a). Following the depletion of 2,3-DCA, *Dehalobacter* continued to grow on 2-CA, with a growth yield of 1.3 ± 0.1 × 10^8^ 16S rRNA gene copies/μmol 2-CA converted (Figure 2b). An extensive review of *Dehalobacter* growth yields measured from various chlorinated organic compounds was conducted previously.^25^ For an easier comparison with other reported values, measured yields from this study are converted into the same units (Table S8). Measured yields from this study are within the range reported previously for 2,3-DCA and 3,4-DCA,^25^ and are close to those previously reported in the literature for other substrates including chlorinated ethanes and benzenes,^25^ falling on the higher end within one order of magnitude. Many studies underestimate the true yield because the impact of cell decay is not considered. This impact becomes more significant when data are analyzed over long-term cultivation (e.g., hundreds of days) and with a limited number of samples. In this study, measurements of cell abundance were frequent and rates were generally faster than many published studies, improving the estimate of yield. Moreover, this study is the first to demonstrate the decay of *Dehalobacter* under stress conditions in the absence of chlorinated organic compounds. After 62 days of starvation, the *Dehalobacter* concentration in the triplicate decay bottles decreased to 35% of its initial level (Figure 2c). The first-order decay constant was calculated as 0.017 ± 0.001 (n=3) day^−1^, corresponding to a half-life of 41 days for *Dehalobacter* without chlorinated electron acceptors. Our results suggest that the common assumption of zero decay in Monod kinetics may not be appropriate for modeling *Dehalobacter* dynamics. Since the decay constant was measured under ample donor conditions, donor limitation in the field could lead to even higher decay rates.

**Figure 2.**
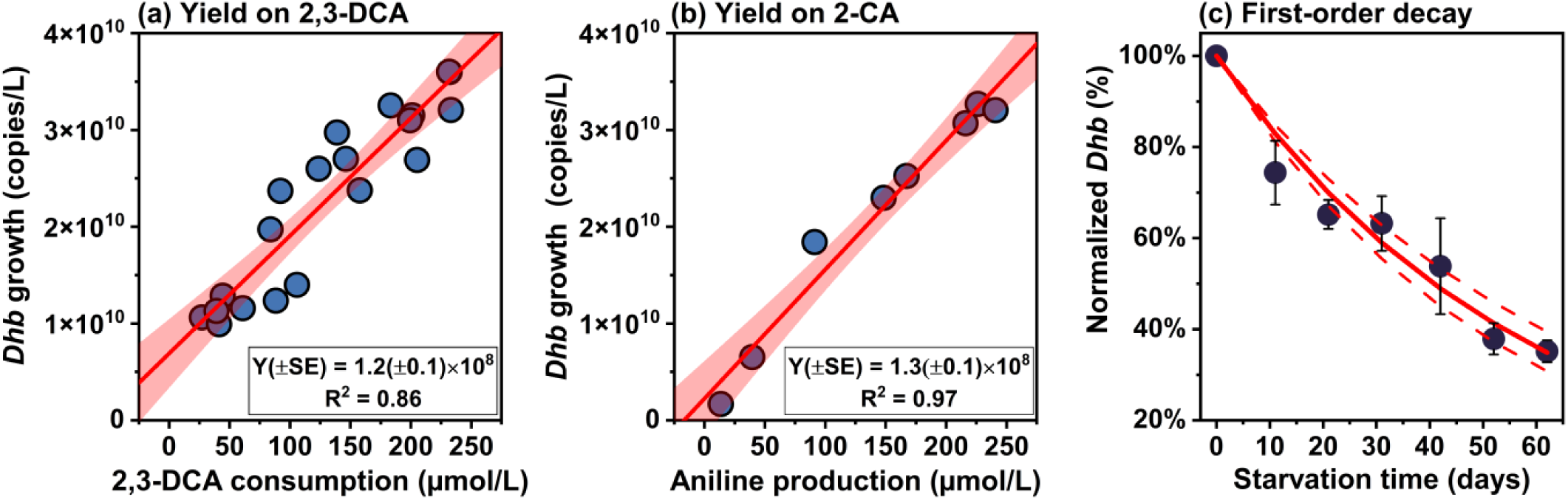
Growth yields (panels a and b) and decay constant (panel c) for *Dehalobacter* (*Dhb*). Growth yields (Y) growing on 2,3-DCA and 2-CA are determined using Ordinary Least Squares regression. Linear regressions are evaluated in Table S10. In panels a and b, solid red lines represent the fitted linear regression, with shaded areas corresponding to the 95% confidence interval. *Dhb* growth denotes the total increase in qPCR-determined *Dehalobacter* absolute abundance at each sampling time relative to the initial abundance at the start of each dechlorination stage. 2,3-DCA consumption represents the cumulative substrate (2,3-DCA) consumption at time t during 2,3-DCA dechlorination, while aniline production indicates the cumulative product (aniline) formation at time t during 2-CA dechlorination. As 2-CA was stoichiometrically converted to aniline, the growth yield on 2-CA was reported in gene copies per μmol 2-CA converted. SE: standard errors associated with the slope. The red line in panel c is the fitted first-order decay model, with dashed lines representing the 95 % confidence interval. Error bars represent the standard deviation in *Dehalobacter* levels normalized to the initial concentration (i.e., C_t_/C_0_) across decay bottles.

**Figure 3.**
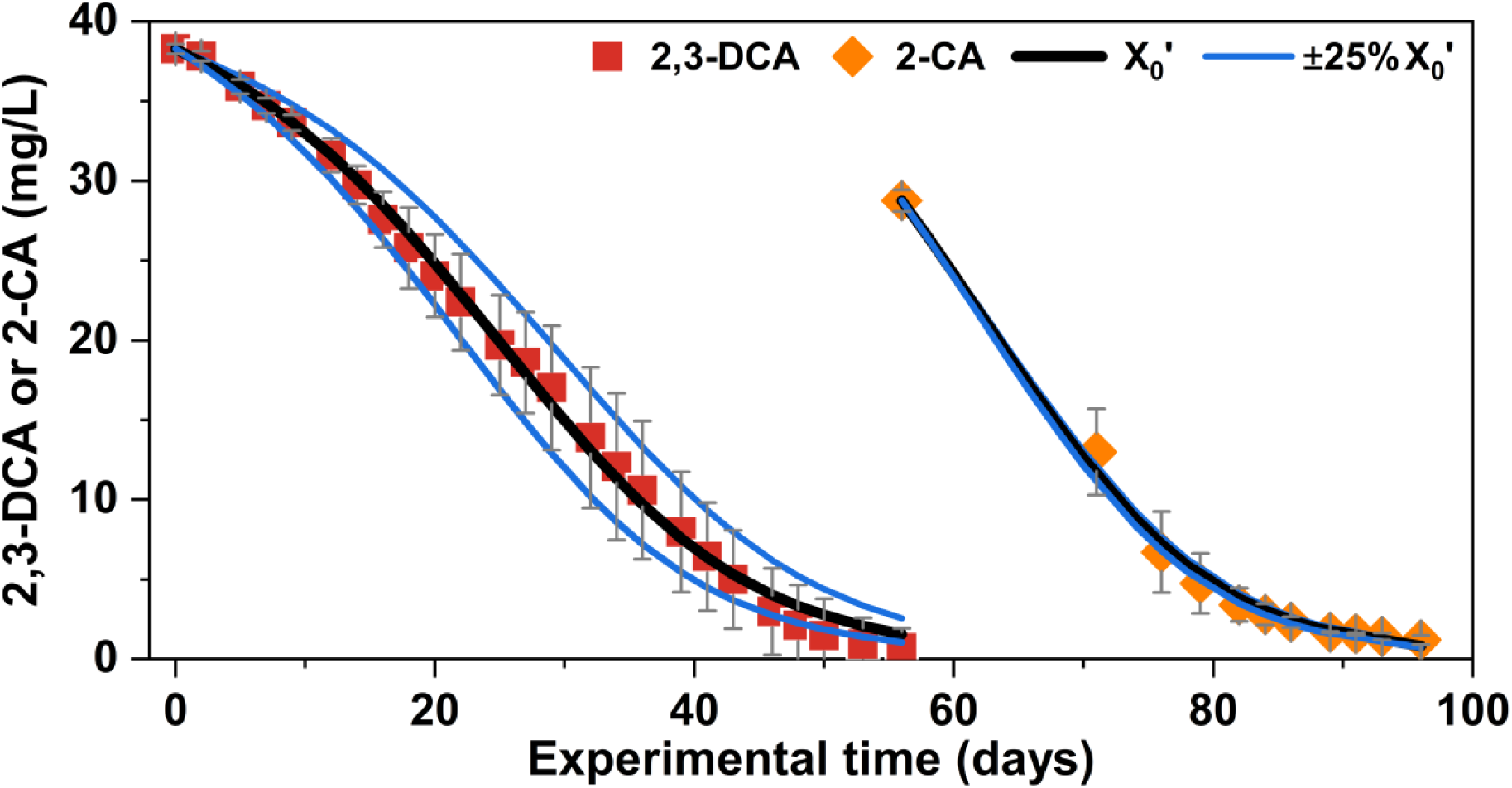
Monod kinetics of dechlorination of 2,3-DCA to aniline. Three simulations were performed at different initial biomass of *Dehalobacter* (75% X_0_^’^, 100% X_0_^’^, and 125% X_0_^’^), using the measured yields and decay constant, along with the optimal values for K_s_ and µ_max_. Red squares and orange diamonds represent the observed depletion data of 2,3-DCA (from all four biotic replicates) and 2-CA (from biotic replicates I and IV), respectively, with error bars showing the standard deviation across the replicates.

We converted the measured yield into mass units (g/g) assuming a mass per cell and 4 16S rRNA gene copies per cell, yielding 1.7 ± 0.2 × 10^−2^ g *Dehalobacter*/ g 2,3-DCA and 2.4 ± 0.2 × 10^−2^ g *Dehalobacter*/ g 2-CA. The theoretical maximum growth yields were also calculated applying the McCarty TEEM model based on microbial energetics,^36,37^ assuming hydrogen as the sole electron donor, ammonium as the sole nitrogen source, and acetate as the sole carbon source (Section S9). The predicted yields at various energy transfer efficiencies, along with the measured yields are compared in Table S9. The thermodynamically predicted yields of *Dehalobacter*, matched the experimental measurements when the energy transfer efficiency was between 45% and 50% (Table S9). The consistency between predicted and measured values gives confidence in the accuracy of the measured yields. The energy transfer efficiency range (45 – 50%) for *Dehalobacter* used in this study are consistent with the reported range (45 – 65%) for anaerobic heterotrophic growth.^37^

#### 6.3.2 Modeling Monod kinetics to estimate K_s_ and µ_max_

With the experimentally determined growth yields and decay constant, kinetics of dechlorination of chloroanilines can be modelled to determine K_s_ and µ_max_ using the Monod equation, assuming electron donor is not limiting. Optimal Monod constants and model evaluation are presented in Table S11. In the case of *Dehalobacter* growth on 2,3-DCA, K_s_ and µ_max_ are determined as 45 ± 16 mg/L and 0.18 ± 0.03 day^−1^, respectively. Due to ample initial electron donor amendment (10-fold the demand for dechlorination of 2,3-DCA to aniline), electron donor limitation unlikely occurred during 2,3-DCA dechlorination. Reproducible trends of 2,3-DCA dechlorination were observed across all four replicates, and the minor variations between replicates can be explained by slight differences (±25%) in the initial biomass (Figure 2). The optimal µ_max_ results in a doubling time for *Dehalobacter* growing on 2,3-DCA without donor limitation ranging from 3.3 to 4.6 days. This range aligns with the doubling time estimated using measured *Dehalobacter* abundance in the initial stage (days 0 to 12), which was 3.3 days. For 2-CA dechlorination, similar constants (K_s_ = 35 ± 24 mg/L and µ_max_ = 0.14 ± 0.06 day^−1^) were found using data from bottles I and IV, exhibiting complete dechlorination. Comparable Monod constants were found for two dechlorination steps. The apparent K_s_ values estimated from the Monod fits are high, likely reflecting not only the intrinsic substrate affinity of *Dehalobacter* but also microbial interactions and possible nutrient limitations within this field-relevant consortium, which can inflate K_s_. Using μ_max_ and the measured K_d_, we estimated the minimum substrate concentration required to support net growth to be approximately 5 mg/L for 2,3-DCA or 2-CA. These values offer more realistic benchmarks for bioaugmentation, though they may vary under different field conditions due to potential changes in K_d._ The µ_max_ of *Dehalobacter* during 2-CA dechlorination was slightly lower than during 2,3-DCA dechlorination, aligning with the thermodynamic predictions that each mole of 2,3-DCA dechlorinated can yield 11% more energy for *Dehalobacter* growth compared to 2-CA, assuming identical energy transfer efficiencies (Table S9). However, kinetics of 2-CA dechlorination varied among individual replicates. Biotic bottles II and III showed that 2-CA dechlorination had slower kinetics, which may indicate that electron donor-limiting conditions were reached during 2-CA dechlorination, as no additional donor was added after T=0.

### 6.4 Roles of other microbes in the consortium

Electron distribution and balances combining results from analyses of chloroanilines, fatty acids, and methane production were investigated to elucidate electron flows and syntrophic relationships within the anaerobic dechlorinating microbial consortium (Figures 4 and S8). This analysis is summarized in Section S10. Other than for biotic replicate IV that exhibited strangely slow acetogenesis, a good electron balance was achieved across replicates, at 84 ± 17 % (Table S12).

**Figure 4.**
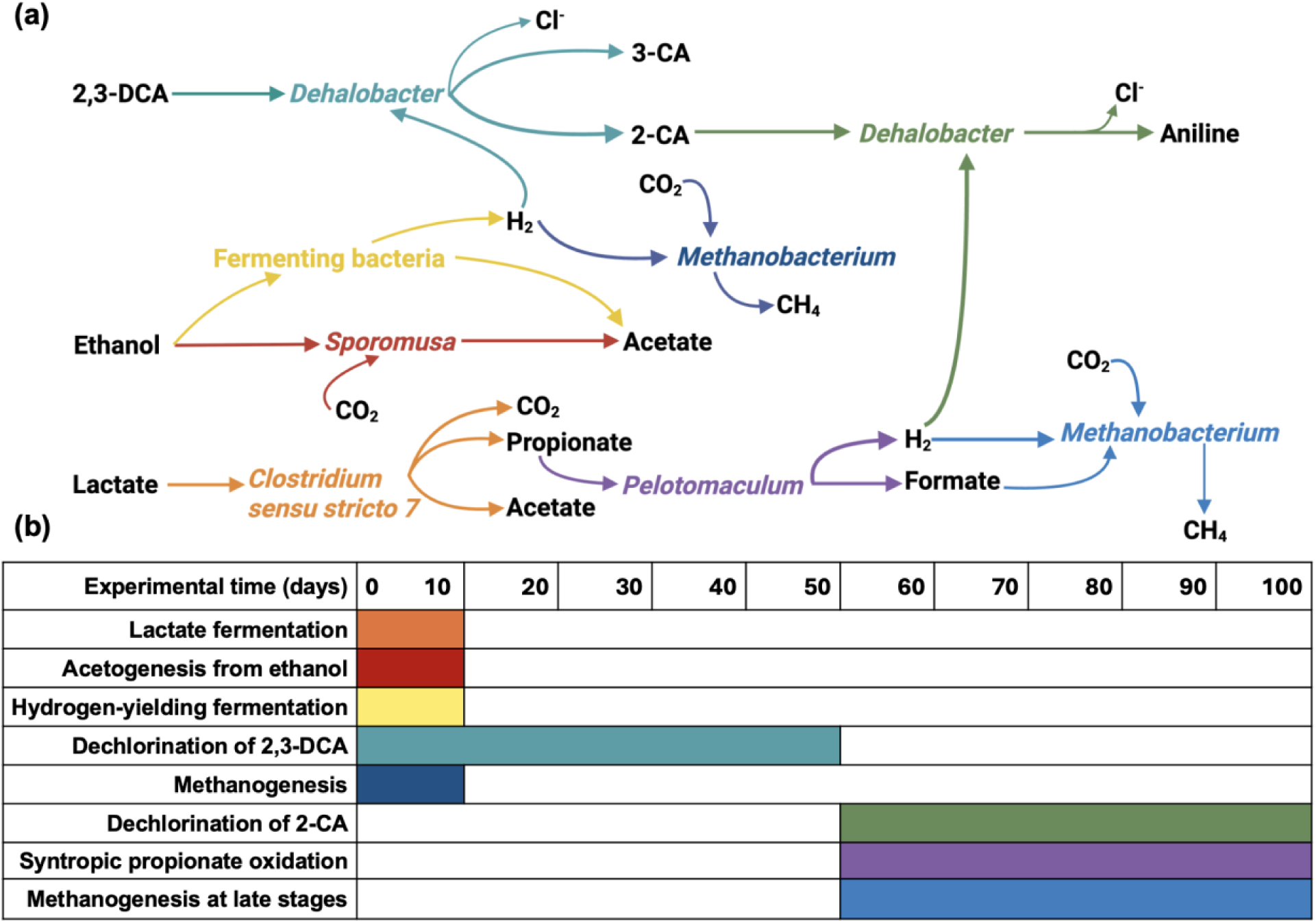
Syntrophic relationships among representative microorganisms in the anaerobic dechlorinating microbial consortium (panel a) and temporal dynamics of major microbiological processes during the experiment (panel b). Electrons available for the whole system originated from the ethanol and lactate. Fermenting bacteria, possibly *Clostridium sensu stricto 7*, *Sporomusa*, and *Sedimentibacter*, consumed ethanol, releasing electrons in the form of hydrogen gas (yellow; the hydrogen-yielding fermentation shown refers to ethanol-based processes), which supported both 2,3-DCA dechlorination (teal) and hydrogenotrophic methanogenesis (navy blue). However, a fraction of ethanol was converted to acetate by *Sporomusa* (red) rather than to hydrogen. In contrast, lactate was not initially used for hydrogen formation; instead, *Clostridium sensu stricto 7* converted most of the amended lactate into acetate and propionate (orange). Subsequently, *Pelotomaculum* oxidized propionate to release hydrogen (purple), supporting 2-CA dechlorination (green) and the late-stage hydrogenotrophic methanogenesis (blue).

#### 6.4.1 Lactate fermentation and acetogenesis from ethanol to produce acetate and hydrogen for *Dehalobacter*

We observed significant changes in the concentrations of volatile fatty acids between days 0 and 5 (Figures 1b and S6), especially production of acetate and propionate from added donors, ethanol and lactate, indicating active fermentative processes early in the experiment. This finding was also supported by 16S rRNA amplicon sequencing and total bacterial concentration data, which revealed a notable growth of all fermenting bacteria, primarily *Clostridium sensu stricto 7* and *Sporomusa*, dominating the bacterial community in the initial stage of the process (Figure 1e and S7a). Additionally, methane production was observed during the same period (between day 0 and day 10, Figure 1d), indicating that methanogenesis aligned with fermentation. Microbial data suggests that fermentative processes were hydrogen-yielding, as the dominant methanogenic genus, *Methanobacterium*, is hydrogenotrophic (Figure S9).

Between days 0 and 5, lactate concentration, initially at 0.6 mM, decreased below the LOQ while acetate, propionate, and formate concentrations increased (Figure 1e). Ethanol initially at 3.3 mM was presumably (the IC used to quantify the VFAs does not detect ethanol) also depleted in the first 5 days because the electron balance is preserved (Figure S8). The stoichiometry ratio of lactate consumed to propionate formed was 2.9 ± 0.2 : 2 (n=4; Section S11) for biotic replicates. This ratio is consistent with the theoretical stoichiometry ratio of lactate: propionate: acetate = 3: 2: 1 during lactate fermentation to propionate and acetate (Section S11).^38,39^ Therefore, this observed ratio confirms that most of the lactate was fermented to propionate and acetate, and that propionate was solely associated with lactate fermentation. The identified *Clostridium sensu stricto 7* was likely responsible for lactate fermentation since species of *Clostridium* were reported to convert lactate to propionate in a ratio of 3:2.^43,44^

Fermentative bacteria can alter their metabolic pathways to produce hydrogen in response to a hydrogen demand.^42^ *Clostridium sensu stricto* has also been documented to produce hydrogen via fermentation.^43–45^ Therefore, *Clostridium sensu stricto 7* along with other identified fermenters, such as *Sedimentibacter* and *Sporomusa*, might be involved in hydrogen-yielding fermentation from ethanol and fatty acids. Partnering with hydrogen consumers such as *Dehalobacter* and methanogens, fermenting bacteria were able to oxidize ethanol to acetate, presumably with hydrogen gas as a product.^46–48^ The electron balance results indicate that the electron demand for complete dechlorination is negligibly low compared to the electrons supplied by ethanol (Figure S8b). Consequently, the hydrogen demand of *Dehalobacter* can be readily met through hydrogen-yielding fermentation of ethanol.

The ratio of initial ethanol addition to measured acetate production, excluding acetate contribution from lactate fermentation, was calculated as 1.2 ± 0.4 (n=4). This stoichiometry is close to fermentation of ethanol to acetate and hydrogen (ethanol: acetate = 1:1)^42^ and acetogenesis from ethanol via the Wood-Ljungdahl pathway (ethanol: acetate = 1:1.5)^49^.

Fermentation reactions are detailed in Section S11. The stoichiometry indicates hydrogen production, and measured products suggest the coexistence of hydrogen-, formate- and non-hydrogen-yielding acetogenesis during fermentation. Acetogenesis was mediated by *Sporomusa*, as indicated by a significant increase in their abundance (Figure 1e). *Sporomusa*, a known homoacetogen, can consume ethanol and CO_2_ to produce acetate through the Wood-Ljungdahl pathway^49^. Furthermore, the electron balance results suggested that only small fraction of electrons from ethanol was used for hydrogen production, as the majority was converted into acetate (Figure S8b).

The acetate pattern in biotic bottle IV differed noticeably from the other replicates, with a slow increase in acetate over the course of the experiment, whereas the other replicates showed an increase mainly within the first 10 days (Figures S6 and S8). This discrepancy on acetate production suggests that biotic bottle IV received an erroneously small amount of ethanol at time zero, potentially due to errors associated with syringe addition. Surprisingly, this discrepancy allows us to identify *Sporomusa* as the primary acetate producer. The limited acetogenesis from ethanol in biotic bottle IV significantly impacted its bacterial composition. In biotic bottle IV, *Sporomusa* was much less abundant in the early stages of the experiment compared to the other replicates, but its abundance gradually increased as the acetate production occurred later (Figure S8). This correlation between acetate production and *Sporomusa* growth suggests that *Sporomusa* is responsible for acetogenesis.

Apart from biotic bottle IV, no notable increase in acetate and propionate was observed after day 10, indicating that lactate fermentation and acetogenesis were much faster processes compared to reductive dechlorination, and were primarily active during the early stages of 2,3-DCA dechlorination (Figure S6). This is further supported by the amplicon sequencing results, which revealed a continuous and significant decline of *Clostridium sensu stricto 7* and *Sporomusa* as the experiment progressed. Beyond day 10, fermentation reactions were much slower owing to lack of readily fermentable substrates. Once amended lactate and ethanol were depleted, the abundance of dominant fermenting bacteria decreased, while *Dehalobacter* became the most abundant bacterial genus (Figures 1e and S7a). Principal component analysis (PCA) of bacterial community composition (detailed in Figure S10) reveals a temporal and reproducible succession under chlorinated aniline stress, with biotic replicates (especially I, II, and III) converging along a shared trajectory. Major community shifts on day 12 (marking the end of fermentation from added donors) and day 50 (the transition between 2,3-DCA and 2-CA dechlorination) observed in the PCA plot (Figure S10) support the proposed temporal dynamics of microbiological processes (Figure 4) and align with VFA and dechlorination profiles.

A variety of fermenting bacteria are involved in interspecies electron transfer within the microbial consortium. They converted fatty acids and alcohols, which *Dehalobacter* cannot utilize for energy and cell synthesis, into hydrogen, formate, and acetate, all of which supported dehalorespiration and growth by *Dehalobacter*. Previous studies have reported *Dehalobacter* relies on hydrogen or formate as the electron donors for anaerobic respiration^50–52^ and acetate as a carbon source.^53^ In this work, acetate uptake into biomass by *Dehalobacter*, estimated from the overall energetic equation (Section S9), was small (< 0.1mM) and not observable amidst the high acetate levels.

#### 6.4.2 Syntrophic propionate oxidation to provide hydrogen and acetate for *Dehalobacter*

During 2,3-DCA dechlorination to monochloroanilines, concentrations of both acetate and propionate, the primary products from fermentation, were steady in the bottles. However, following the depletion of 2,3-DCA and initiation of dechlorination of 2-CA to aniline, propionate levels began to decrease (days 50 – 95), while acetate concentrations exhibited minimal decline (Figure 1b). A moderate amount of methane was also produced over the same period, following a phase of negligible methanogenesis throughout 2,3-DCA dechlorination (Figure 1d). This late-stage methanogenesis can be attributed to the decrease in propionate concentrations (biotic bottles I, II, and IV), with a recovery of 78 (± 11) % (n=3), calculated as the electron equivalent ratio of the methane production to the reduction in propionate. Moreover, this late-stage methanogenesis was associated with a rise in the bacterial abundance of *Pelotomaculum* (Figure 1e). These are syntrophic propionate-oxidizing bacteria that are obligately co-cocultured with methanogens.^54,55^ Thus experimental evidence indicates that propionate was consumed by *Pelotomaculum* via syntrophic oxidation. Subsequently, the products from syntrophic propionate oxidation, formate and hydrogen, were used by hydrogenotrophic methanogens to form methane and by *Dehalobacter* for 2-CA dechlorination. The syntrophic oxidation of propionate had an important role in this anaerobic dechlorinating system. Lactate fermentation observed in the initial stage of the experiment produced propionate and acetate, which was not a hydrogen-yielding process. In addition, propionate cannot be directly utilized by *Dehalobacter* or methanogens and therefore built up (days 10 – 50). Thanks to *Pelotomaculum*, propionate was metabolized into formate and hydrogen, thus enhancing the electron transfer efficiency from the amended lactate to hydrogen gas and keeping the propionate concentration low.

Additionally, our results clearly demonstrate that syntrophic propionate oxidation and associated hydrogenotrophic methanogenesis did not commence until the depletion of 2,3-DCA (Figure 1). This relatively late onset of syntrophic propionate oxidation may result from the toxicity of 2,3-DCA on *Pelotomaculum* or methanogens. Syntrophic propionate oxidation is energetically unfavourable unless coupled with effective hydrogen-consuming processes (e.g., dechlorination and methanogenesis) to maintain low hydrogen patrial pressures.^47^ During 2,3-DCA dechlorination, *Dehalobacter* likely outcompeted methanogens for hydrogen, as shown by the consistent and reproducible dechlorination profiles, while methane production remained minimal. However, the hydrogen demand for dechlorination, as previously discussed, was too low to trigger the syntrophic propionate oxidation.

#### 6.4.3 Acetate accumulation and potential inhibition of chloroanilines on acetoclastic methanogens

The accumulation of acetate in the dechlorinating cultures was unexpected, but also observed in our companion study^25^. Rather than methane, most of electrons from added lactate and ethanol ended up in the form of acetate (Figure S8). The acetate accumulation suggests a lack of effective acetate consumer in the community, which is further confirmed by the 16S rRNA sequencing results (Figure S9). All identified methanogens, such as *Methanobacterium*, which dominated the Archaea community with a relative abundance greater than 97%, and *Methanoregula*, are hydrogenotrophic methanogens that produce methane exclusively from hydrogen and CO₂ and not from acetate.^56,57^ Thus, acetate accumulation was due to the absence of acetoclastic methanogens. The absence of acetoclastic methanogens suggests they were eliminated by elevated concentrations of chlorinated anilines, and effectively diluted out of the culture after successive transfers. Previous studies have reported that chlorinated compounds and amino-substituted aromatics can inhibit conversion of acetate to methane.^58,59^ Similar acetate accumulation under stress conditions was reported in tetrachloroethene-dechlorinating microcosms exposed to high salinity, where upstream fermentation (acidogenesis and hydrogenesis) remained active, but downstream consumers of fermentation products were inhibited.^60^ Excessive acetate not only led to a low electron transfer efficiency from the amended electron donors to hydrogen gas but may also exhaust the buffering capacity of the mineral medium, leading to low pH conditions that could harm the microbial system. Thus, for dechlorinating microbial systems lacking the acetoclastic methanogens, regular culture transfer can be a preventive measure against the excessive acetate. Moreover, acetate accumulation can serve as a potential indicator of methanogenesis inhibition in the groundwater.

## 7 Environmental Implications

This study presents the stoichiometric reductive dechlorination of 2,3-DCA by *Dehalobacter*, resulting primarily in aniline via 2-CA, but also in the low (<15%) yet persistent accumulation of 3-CA. This underscores the importance of monitoring transformation products during dechlorination processes, which is analogous to the accumulation of vinyl chloride during dichloroelimination of 1,2-diochloroethane to ethene.^61^ Thanks to a unique high-frequency sampling regime, we estimated kinetic parameters and growth yields that are valuable for modeling the fate and transport of chloroanilines in groundwater and to quantify *in situ* dechlorination. The syntrophic interactions between *Dehalobacter* and its microbial community can be used as one line of evidence to identify natural attenuation of chlorinated anilines at contaminated sites. Co-contaminants of chloroanilines at the site, primarily chlorobenzenes and chloronitrobenzenes, likely exert selective pressure by favoring organohalide-respiring bacteria such as *Dehalobacter*. *Dehalobacter* with broader dechlorination capabilities may have been enriched through dechlorination of chlorobenzenes and chloroanilines that are also formed via nitro-group reduction of chloronitrobenzenes, as observed in our companion study^25^ using cultures derived from the same site. In addition, the achievement of complete dechlorination at a high substrate level (40 mg/L) with a low inoculum (3% v/v) holds promise for the application of biostimulation (with electron donors) and bioaugmentation strategies to address chloroaniline contamination. This study also explored interspecies electron transfer as a function of amended donors over time, deciphering some of the dynamics of successive microbial processes, including the issue of acetate accumulation, offering insights into establishing and maintaining a stable anaerobic dechlorinating system in culture or *in situ*. Although hydrogen was not directly quantified in this work, it is presumed to be the electron donor based on the observed fermentation pathways, electron balance, and the established physiology of *Dehalobacter*. While this work prioritized a mixed, field-derived consortium to reflect the site conditions, future studies should directly measure hydrogen and use it as the sole electron donor to confirm its role. Our ongoing work involves metagenomics and compound-specific isotope analysis to elucidate the reaction pathways of chlorine removal and determine isotope enrichment factors for field-based assessment of *in situ* dechlorination of chloroanilines.

## 8 ASSOCIATED CONTENT

The supporting information is available free of charge. Supplementary descriptions of subculture establishment, analytical methods, and data analysis, along with the relevant figures and tables discussed in the paper (PDF). Original data provided in spreadsheet format (.xlsx).

## 9 ACKNOWLEDGMENTS

This research was supported by the Natural Sciences and Engineering Research Council of Canada (NSERC) Discovery (RGPIN-2015-06460), Alliance (ALLRP 575243-22), and Canada Research Chair (950-230892) grants to E. Passeport, and a Mitacs (IT29192) grant to both E. Passeport and E. Edwards. The authors declare no conflicts of interest. This publication is in accordance with the Brazilian biodiversity law n.° 13.123, from May 20th, 2015. This is non-profit research and has no commercial intention between the parties. This is a collaboration project between the parties involved. Access Registration No. A3D9F11.

## Notes

### Competing Interest Statement

The authors have declared no competing interest.

### Summary of Updates

No changes made to the manuscript Only supplemental files updated.

