## Supporting information for "Reductive dechlorination of 2,3-dichloroaniline by *Dehalobacter* in an anaerobic enrichment culture"

### Table of Contents

#### 1. List of sections

**Section S1. Dechlorination profiles, cultivation conditions, and maintenance protocols for the parent and transfer cultures .......... S5**

**Section S2. Details of enrichment culture setup and feeding procedures .......... S10**

**Section S3. HPLC/DAD analysis .......... S11**

**Section S4. GC/FID analysis .......... S12**

**Section S5. IC analysis .......... S12**

**Section S6. Sample collection for DNA extractions .......... S13**

**Section S7. qPCR operating protocols .......... S14**

**Section S8. Processing of 16S rRNA amplicon sequencing results .......... S15**

**Section S9. Calculation of theoretical yields for Dehalobacter on 2,3-DCA and 2-CA using the TEEM method .......... S15**

**Section S10. Analysis of electron distribution and balances .......... S19**

**Section S11. Stoichiometry of electron donor conversion during fermentation .......... S20**

#### 2. List of figures

**Figure S1. Origin of the subcultures (WANG-23DCA-T1, WANG-23DCA-T2, and WANG-23DCA-T3) and biotic bottles in this study .......... S6**

**Figure S2. Dechlorination profile of enrichment culture WANG-23DCA-T1. .......... S7**

**Figure S3. Dechlorination profile of enrichment culture WANG-23DCA-T2. .......... S8**

**Figure S4. Dechlorination profile of enrichment culture WANG-23DCA-T3. .................. S10**

**Figure S5. Dechlorination of 2,3-DCA to aniline in four biotic replicates .......................... S22**

**Figure S6. Concentrations (mmol/L) of lactate, acetate, propionate, and formate in four biotic replicates (bottles I to IV). ........................................................................... S23**

**Figure S7a. Relative abundance of bacteria at the genus level (determined from amplicon sequencing) combined with the absolute abundance of total bacteria (quantified by qPCR targeting general bacteria, reported as 16S rRNA gene copies/L) for all biotic bottles (I to IV) during the course of the experiment. .................................................................. S24**

**Figure S7b. Initial bacterial compositions along with the absolute abundance of total bacteria (16S rRNA gene copies/L) for all four biotic replicates (bottles I to IV). ............ S25**

**Figure S8. Bacterial composition (panel a) alongside electron balances and distributions (panel b) for all four biotic replicates (I to IV) during dechlorination of 2,3-DCA to aniline. ........................................................................................................................ S26**

**Figure S9. Relative abundance of archaea at the genus level in biotic bottles (I to IV) during the course of the experiment. ........................................................................ S28**

**Figure S10. (a) Principal component analysis (PCA) of bacterial community composition across biotic bottles during the incubation period. (b) Heatmap of relative abundances of main bacterial genera over time……….……………………………………………………S29**

#### 3. List of tables

**Table S1. qPCR raw data and calculations (.xlsx file)**

**Table S2. 16S rRNA gene sequence, taxonomic assignment, and relative abundance of bacterial ASVs identified in DNA samples using Illumina sequencing (.xlsx file)**

**Table S3. 16S rRNA gene sequence, taxonomic assignment, and relative abundance of archaeal ASVs identified in DNA samples using Illumina sequencing (.xlsx file)**

**Table S4. Sequence alignments for predominant *Dehalobacter* ASVs involved in chloroaniline dechlorination…………………………………………………………..........S31**

**Table S5. Data for chlorinated amino aromatics analysis…………………………………S32**

**Table S6. Data for analysis of volatile fatty acids and inorganic anions…………………..S33**

**Table S7. Data for headspace methane analysis and calculated methane production…....S35**

**Table S8. Summary of measured growth yields of *Dehalobacter* (Dhb) in various units...S35**

**Table S9. Comparison of measured and predicted growth yields for *Dehalobacter* (Dhb) at different values of the energy transfer efficiency (ɛ) ………………………………………S36**

**Table S10. Evaluation of linear regressions for calculating the growth yields of *Dehalobacter* associated with 2,3-DCA dechlorination (days 0 to 50) and 2-CA dechlorination (days 50 to 100)……………………………………………………………S37**

**Table S11. Results and evaluation of the Monod kinetics to determine Monod constants.S37**

**Table S12. Calculation of electron balances and distribution…………………………...S38**

#### Section S1. Dechlorination profiles, cultivation conditions, and maintenance protocols for the parent and transfer cultures

On April 21st, 2022, a 10 mL subsample was taken from enrichment culture 23DCA-T2[1] established and maintained by Araújo et al.(2025). Until May 4, 2022, the subsample was stored in the anaerobic glove box, which is described in the main text. On May 4, 2022, this subsample was used as inoculum for a set of new sub-cultures in preparation for kinetic and stable isotope analyses. The first new subculture, WANG-23DCA-T1, was performed by inoculating 90 mL of anaerobic medium with the original 10 mL sample. Culture WANG-23DCA-T1 was repeatedly amended with 2,3-DCA and electron donors, lactate and ethanol. Once dechlorination and growth were observed, the culture was topped up with medium multiple times to build up culture volume for future experiments. These transfers (WANG-23DCA-T1, T2 and T3) are described below (Figure S1-S4). Before the subsample was taken, the parent enrichment culture (23DCA-T2) had been fed with 2,3-DCA at 10 mg/L (62 µM) for several hundred days and regularly converted 2,3-DCA to 3-CA. To try to increase the dechlorination rate, culture WANG-23DCA-T1 was re-spiked with double the 2,3-DCA concentration, i.e., 20 mg/L (125 µM). With each 2,3-DCA addition (20 mg/L), lactate and ethanol were spiked at 0.5 mM and 2 mM in the enrichment culture, respectively. The pre-reduced anaerobic mineral medium used for WANG-23DCA-T1 was as described previously[2] except that ammonium was omitted – originally to encourage microbes that could use chloroanilines as the source of nitrogen.

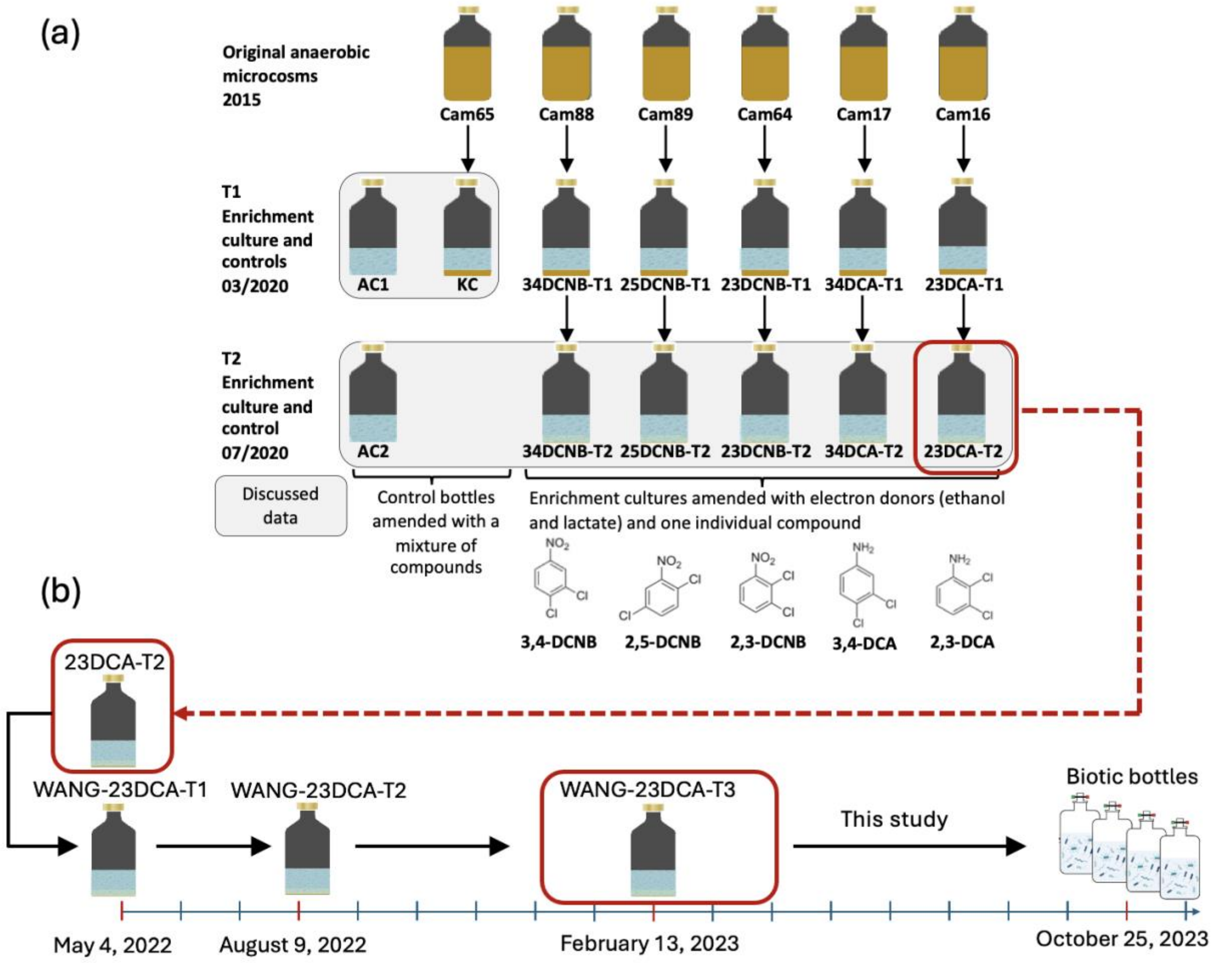


**Figure S1. Origin of the subcultures (WANG-23DCA-T1, WANG-23DCA-T2, and WANG-23DCA-T3) and biotic bottles in this study. (a) History of the preceding microcosms and enrichment cultures. Culture 23DCA-T2[1] was used to inoculate the first subculture (WANG-23DCA-T1). (b) Timeline of the establishment of the subcultures. Panel (a) was adapted from Figure S1 of the companion work[1].**

The dechlorination curve of WANG-23DCA-T1 is shown in Figure S2. Day 0 of Wang-23DCA-T1 is May 4, 2022 (Figure S2). For the first 97 days, 2,3-DCA dechlorination to 2-CA was slow (< 0.1 mg/L per day). 2-CA persisted as the major end product in the bottle (Figure S2). To try to enhance the transformation rate of 2,3-DCA, we decided to add back ammonium to the concentration in the original medium ($C_{NH4+}$=10 mM). After day 97, culture WANG-23DCA-T1 was maintained in the regular ammonium-containing medium, and electron donor additions increased to 0.6 mM for lactate and 3 mM for ethanol, while the 2,3-DCA feeding concentration remained unchanged at 20 mg/L. After switching to the ammonium-containing medium, the rate of dechlorination gradually increased, from feeding once every 50 days to once every two weeks (Figure S2). By around day 300, trace amounts of aniline were observed at low µM levels. Killed controls with autoclaved enrichment culture were performed in the companion work[1], showing no degradation or transformation activity.

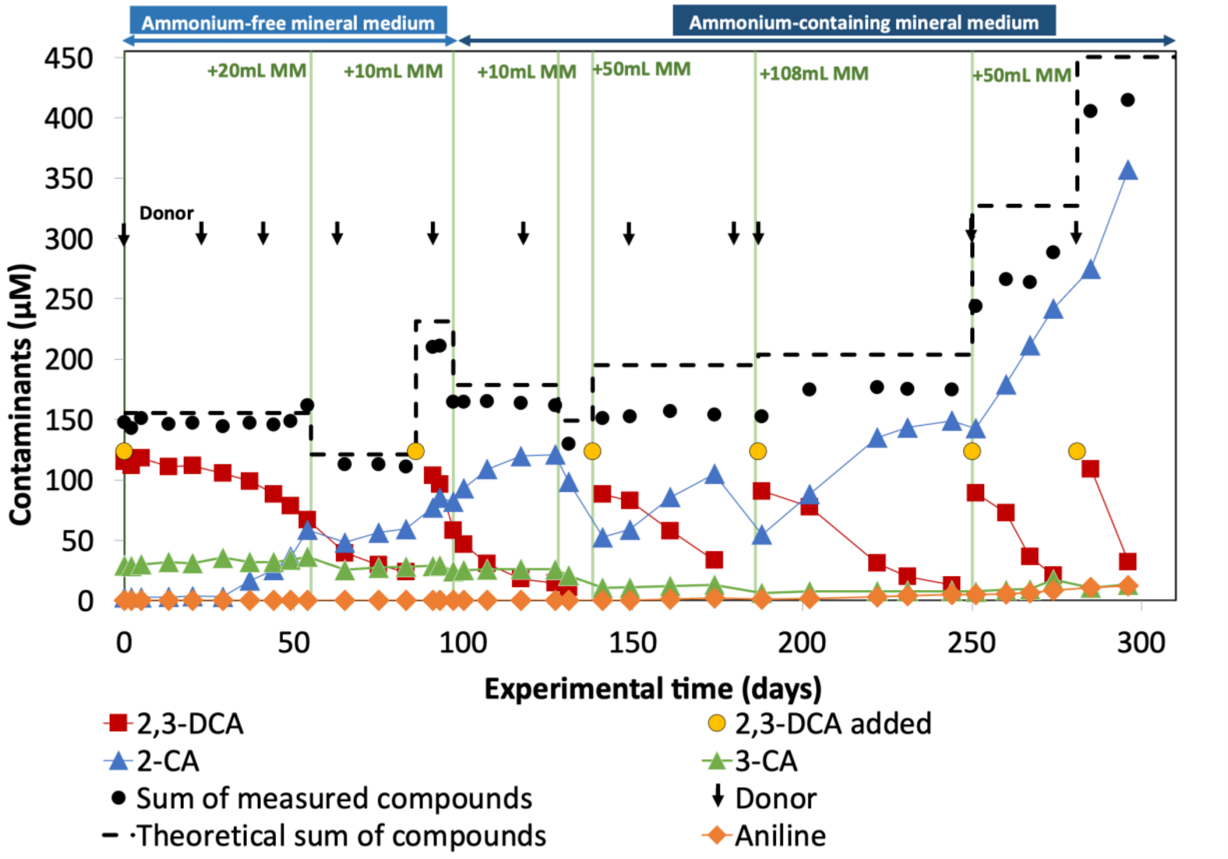


**Figure S2. Dechlorination profile of enrichment culture WANG-23DCA-T1. Arrows indicate time points for electron donor addition. Green vertical lines show time when additional medium was added to increase the volume of culture. The initial volume was 100 mL, and the final volume of the culture after 300 days was around 220 mL (the sample volume for concentration analysis was around 1.5 mL when growing these subcultures). Day 0 of Wang-23DCA-T1 is May 4, 2022. A sample of 50 mL was withdrawn on day 97 (August 9, 2022) for inoculating WANG-23DCA-T2. After day 97,**

**culture WANG-23DCA-T1 was still maintained in parallel**

On day 97 (August 9, 2022), a 50 mL portion of culture WANG-23DCA-T1 was transferred to a new bottle to establish a new enrichment culture, designated WANG-23DCA-T2. To this initial 50 mL culture, 0.5 mL of 1000 mM ammonium chloride was added, to bring the ammonium concentration to 10 mM. Culture WANG-23DCA-T2 was scaled up from 50 mL to 210 mL using the ammonium-containing medium, with maintenance protocols remaining unchanged (Figure S3). This second transfer continued to produce mostly 2-CA from 2,3-DCA with traces of aniline. However, the transformation rate significantly increased over time, requiring a more frequent feeding (from approximately once per 50 days to once per 25 days).

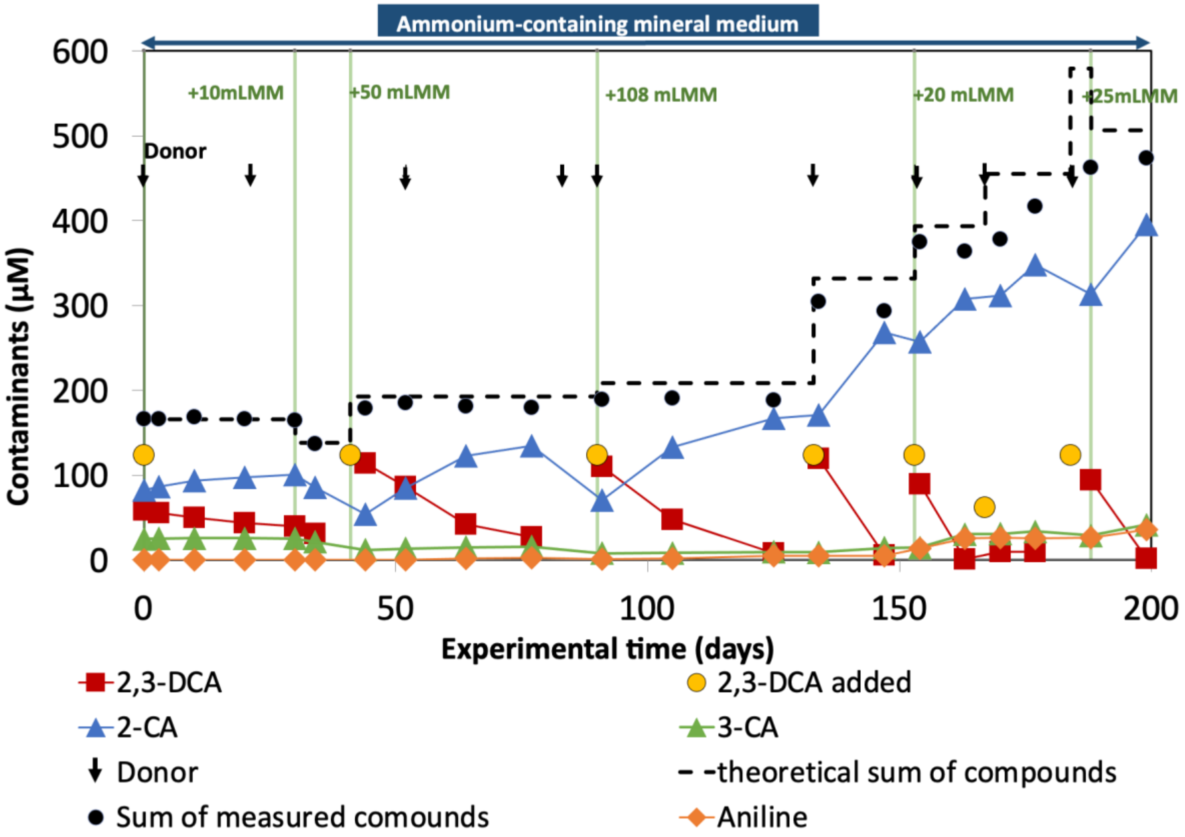


**Figure S3. Dechlorination profile of enrichment culture WANG-23DCA-T2. Day 0 of WANG-23DCA-T2 is August 9, 2022. After six rounds of feeding 2,3-DCA at 20 mg/L, the inoculation for setting up subculture WANG-23DCA-T3 took place on day 188.**

On day 188 of WANG-23DCA-T2 (February 13, 2023), a 12 mL portion of culture from WANG-23DCA-T2 was transferred into 388 mL of regular medium (a 3% transfer) to establish culture WANG-23DCA-T3 (400 mL). Fresh medium was periodically added to this culture to build up a volume of ~ 1 L. After day 147 (July 10, 2023) of this third transfer and medium top-ups, aniline formation dramatically increased, resulting in complete conversion of accumulated 2-CA to aniline (Figure S4). This third transfer was the inoculum of the experiments described herein. The inoculum for this study was collected on day 254 of WANG-23DCA-T3 (October 25, 2023).

In the culture WANG-23DCA-T3, 2,3-DCA was typically fully converted within two weeks after each feeding. Interestingly, 2-CA was no longer the only end-product. Instead, more significant accumulation of aniline was observed. The conversion of 2-CA to aniline became most pronounced after day 147, when 2-CA concentrations exceeded 1000 μM. In contrast, 2,3-DCA concentrations remained low during this period because of the rapid rate of microbiological consumption, preventing the accumulation of 2,3-DCA in the system. Therefore, we hypothesize that 2,3-DCA can undergo reductive dechlorination to 2-CA, which can be further dechlorinated to aniline if 2,3-DCA is low in the system. However, due to limited chemical and microbiological evidence, infrequent sampling, and poor mole balance during long-term cultivation (possibly due to inconsistent 2,3-DCA feeding amounts), the reaction pathway, responsible bacteria, and stoichiometry remained unclear, prompting this study.

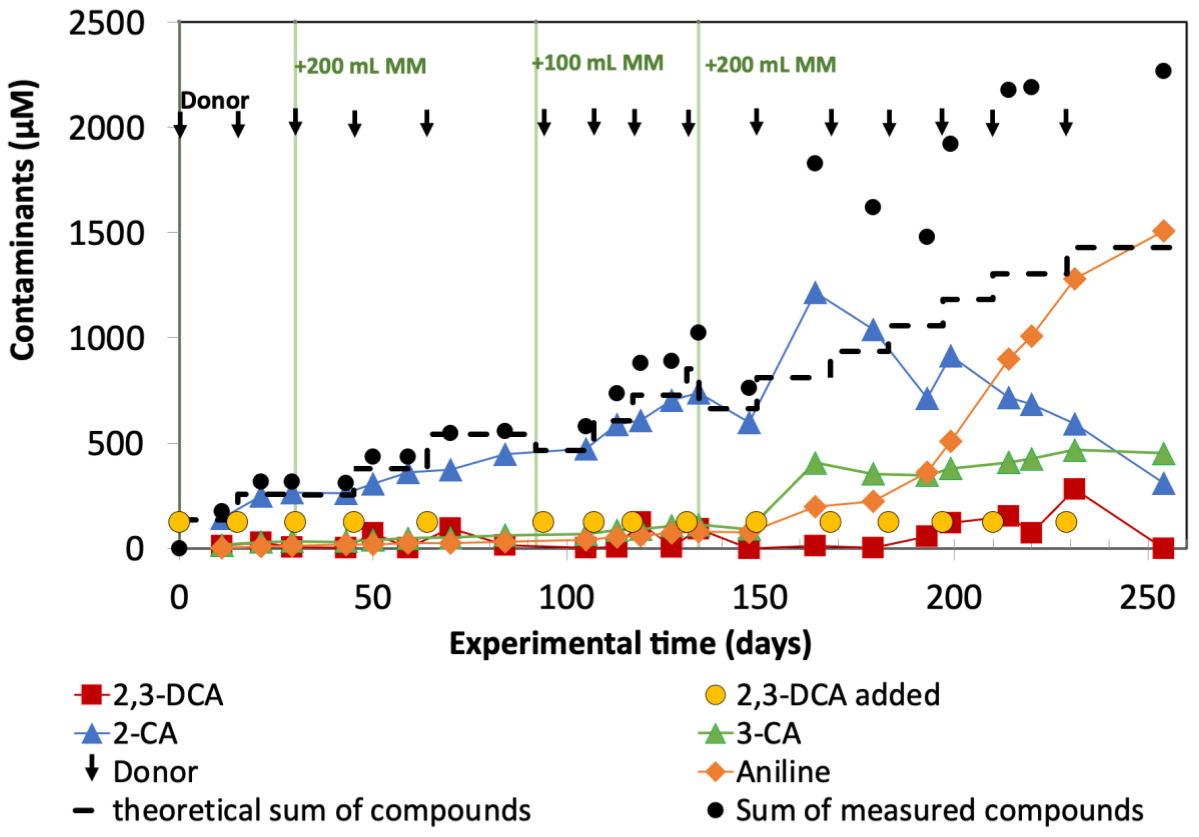


**Figure S4. Dechlorination profile of enrichment culture WANG-23DCA-T3. Day 0 of Wang-23DCA-T3 is February 13, 2023. The inoculum for this study was taken on day 254 (October 25, 2023).**

**Section S2. Details of enrichment culture setup and feeding procedures**

All 250-mL Boston round glass bottles were sterilized by autoclaving at 121 °C and 15 psi for 30 minutes. The Mininert valves with PTFE liners were sterilized by wiping with alcohol pads. Prior to setting up the bottles, autoclaved 250-mL Boston round glass bottles, Mininert valves, and consumables, such as sterile needles, syringes, and serological pipettes, were placed inside the glove box and then stored for a minimum of two days to equilibrate to anaerobic conditions. The feeding vials were prepared as follows. A single-analyte feeding stock solution for 2,3-DCA was prepared in acetone at a concentration of 10 g/L. For a target substrate concentration at 40 mg/L in experimental bottles, 840 µL of the 10 g/L stock solution was added to each sterilized 2 mL HPLC glass vial. To avoid introducing acetone into the enrichment cultures, feeding vials were then left loosely capped and placed in a fume hood to ensure complete acetone evaporation. Once acetone was completely evaporated, the feeding vials were taken into the glove box and sat with loose caps for 2 days to remove residual oxygen prior to being capped until use.

**Section S3. HPLC/DAD analysis**

Each sampling involved collecting 350 µL of liquid from the enrichment cultures using a 2.5 mL Hamilton gastight glass syringe with a PTFE Luer Lock, followed by filtration through a 0.22-µm hydrophilic PTFE syringe filter. Filtered samples were stored at 4 °C before analysis. Concentration analysis was done immediately after sampling. The HPLC column oven was maintained at 30 °C. A gradient elution profile was run starting from 30% acetonitrile and 70% water for 3.5 minutes, ramping up to 75% acetonitrile and 25% water in 1 minute, and a final hold for 3.5 minutes, prior to returning to initial conditions held for 8 minutes. The constant flow rate was 0.3 mL/min. Samples were introduced using the built-in autosampler. Quantification was performed at 254 nm. Mixed standard solutions with known concentrations for HPLC/DAD calibration were prepared with compounds in Milli-Q water and analyzed together with samples for every analysis sequence under similar instrument conditions. The concentrations of compounds in unknown samples were determined using calibration curves that were generated from the analysis of external standard solutions ($R^2 > 0.99$ and p-value $< 3 \times 10^{-7}$ for all target compounds). Six external calibration standards were prepared in the range of 0.1 mg/L to 50 mg/L. The vessels storing calibration standard solutions, wrapped with aluminum foils to prevent photolysis, were kept at 4 °C for long-term storage (less than 3 weeks). Repeatability (intra-run precision, expressed as the relative standard deviation among replicate measurements within a day) of less than 0.5% (n=7) was observed for all compounds above 1 mg/L. Reproducibility (inter-run precision, expressed as the relative standard deviation among measurements taken over seven days within a month) of less than 2% (n=7) was observed for all compounds above 1 mg/L.

**Section S4. GC/FID analysis**

In this work, GC/FID was used to quantify methane production. For this purpose, headspace samples were obtained roughly every ten days. Collection of headspace samples, each measuring 0.3 mL, was performed using a gas-tight glass syringe with an on-off button valve and a volume capacity of 0.5 mL. Collected samples were immediately introduced into the GC equipped with a GSQ I.D. PLOT column (30 m × 0.53 mm) manufactured by J & W Scientific (Folsom, CA, USA). The carrier gas employed was helium, and the column head pressure was consistently maintained within the range of 22-24 psi. The column oven temperature was constant at 190 °C, while the injection port and FID temperatures were controlled at 200 °C and 250 °C, respectively. Calibration was conducted using external gas standards ($R^2 > 0.99$). Benzene was not detected in any of the samples. Methane production was calculated by combining the measured headspace methane with the methane dissolved in the liquid phase, which was estimated using Henry's law and a Henry's constant at $1.4\times10^{-5}$ mol/($m^3\cdot$Pa).[3]

**Section S5. IC analysis**

Collected IC samples underwent syringe filtration through a 0.22-μm nylon syringe filter. The filtered samples were collected in 1.5 mL Eppendorf tubes and stored at -20 °C until further processing and analysis. Prior to analysis, all samples were thawed at 4°C and then vortexed for 5 seconds. The well-mixed thawed samples underwent a 1:10 dilution using Milli-Q water, with 110 μL of the samples being added to 990 μL of Milli-Q water in 2 mL Eppendorf tubes. Subsequently, the diluted samples were vortexed for an additional 5 seconds.

Separation of anions and fatty acids was achieved at a constant flow rate of 0.25 mL/min using a Dionex™ Integrion IC system with a Dionex™ IonPac™ 2 × 250 mm AS11-HC column. The injection volume was set at 450 µL. The instrument was equipped with an automated eluent generator and an eluent cartridge (Dionex™ EGC 400 KOH), and the following eluent generation method was used: potassium hydroxide gradually increased from 0.5 mM to 2.5 mM during the first 10 minutes, then ramped up to 30 mM from 10 to 29 min, where it was maintained at 30 mM until 40 min. It was subsequently reduced to 0.5 mM from 40 to 42.1 min and held steady at 0.5 mM up to 47 min. External calibration standards were prepared, combining all target anions and volatile fatty acids, at concentrations of 2, 1, 0.5, 0.2, 0.05, 0.01, 0.005, and 0.001 mM ($R^2 > 0.99$ for all target anions and volatile fatty acids). LOQs for target anions were calculated using the following equation:

$$LOQ = \frac{10 \times background}{s}$$

where background is the average peak area of the target anion measured in the technical blanks (n=7) and s is the slope of the calibration curve for the target compound. If the calculated LOQ is less than the lowest level of the external standards (i.e., 0.001 mM), we reported 0.001 mM as the LOQ.

**Section S6. Sample collection for DNA extractions**

Enrichment cultures were shaken by hand for 30 seconds before sampling. Each collected cell sample (4 mL) was evenly split into two 2-mL sterile microcentrifuge tubes (tube A and tube B), followed by centrifugation at 13,000 rpm for 20 min at 4 °C. After centrifugation, 1.75 mL of the supernatant was removed from tube A using pipettes with sterile tips. In tube B, 1.5 mL of the supernatant was discarded, and the cell pellets were resuspended in the remaining liquid. This resuspended sample (500 µL) was then transferred into tube A. Tube A, containing a 750-µL sample combining cells from both tubes, underwent a secondary round of centrifugation at 13,000 rpm for 20 min at 4°C. Subsequently, 550 μL of the supernatant was carefully eliminated, leaving only 200 μL of cell pellets retained in the tube.

**Section S7. qPCR operating protocols**

All DNA extracts for qPCR analysis were diluted at least 10 times using DNA-free UltraPure DNase/RNase-free distilled water (Invitrogen, Carlsbad, CA) to minimize matrix effects that could impact qPCR reactions. A standard curve was generated through a dilution series ranging from 10 to $10^7$ copies/μL. This series was prepared by diluting plasmid (i.e., *Dehalococcoides* and *Dehalobacter* plasmids used for quantification of total bacteria and *Dehalobacter*, respectively) with a known absolute abundance in UltraPure water (Table S1). In a UV-treated PCR cabinet, a Master Mix was prepared in 0.5-mL DNA LoBind tubes (Eppendorf Canada), combining 10 μL of SsoFast EvaGreen Supermix (Bio-Rad Laboratories, Mississauga, ON), 6 μL of UV-treated UltraPure water, and 1 μL each of 10 μM forward and reverse primers. For each qPCR reaction, 18 μL of the Master Mix was mixed with 2 μL of either DNA samples or plasmid dilutions. Triplicate reactions were set up for each plasmid dilution, while duplicate reactions were prepared for each DNA sample. Technical blanks, consisting solely of 18 μL of the Master Mix, were also included in the analysis. qPCR reactions were conducted on a C1000 Touch thermal cycler coupled with CFX96 Touch real-time PCR detection system (Bio-Rad Laboratories, Hercules, CA). Thermocycling process included an initial denaturation step at 98°C for 120 s, followed by 40 thermal cycles, and an extension at 65°C for 5 s. Each thermal cycle involved denaturation at 98°C for 5 s and annealing at a lower temperature for 10 s (62.5°C for *Dehalobacter* and 55°C for general bacteria). qPCR results were analyzed using Bio-Rad CFX Manager software.

#### Section S8. Processing of 16S rRNA amplicon sequencing results

The 16S rRNA amplicon sequencing results were processed using Quantitative Insight Into Microbial Ecology (QIIME 2 v.2021.2). Original amplicon reads underwent trimming to remove primer sequences and staggered ends, followed by truncation and denoising using the DADA2 plugin in QIIME 2. Forward reads of 260 bp and reverse reads of 240 bp were quality-filtered and then merged, permitting up to 2 expected errors in the overlapping region. Sequences identified as chimeric or shorter than 400 nucleotides were discarded. The final amplicon sequence variants (ASVs) were classified using the SSU SILVA v.138 database. To confirm accuracy, the taxonomy of the most abundant ASVs used in graphing was cross-checked against the NCBI database.

#### Section S9. Calculation of theoretical yields for *Dehalobacter* on 2,3-DCA and 2-CA using the TEEM method[4]

All $\Delta G^{0\prime}$ values used in the following energetics calculation represent the Gibbs free energy change of reactions under standard conditions (i.e., 25°C, 1 atm, and an activity of 1 for all reactants and products present in the reaction) and at a pH of 7.0. For the energetics calculation, hydrogen was assumed to be the electron donor and chloroanilines (2,3-DCA and 2-CA) were electron acceptors. Also, ammonium was the nitrogen source, and acetate was assumed to be the carbon source for cell synthesis.

Calculation of theoretical yield for *Dehalobacter* on 2.3-DCA:

**i.** With hydrogen as the electron donor, the electron donor half-reaction ($R_d$) is

$$R_d: \quad H^+ + e^- \rightarrow \frac{1}{2}H_2 \qquad \Delta G_d^{o\prime} = 39.87\ kJ/e^-eq$$

The $\Delta G_d^{o\prime}$ value was obtained from the textbook 'Environmental Biotechnology: Principles and Applications' authored by Rittmann and McCarty (2001).[5] Unless specified otherwise, all the free energy values mentioned in the following text were obtained from this textbook.

**ii.** With 2,3-DCA as the electron acceptor, the electron acceptor half-reaction ($R_a$) is

$$R_a: \quad \frac{1}{2}C_6H_3Cl_2NH_2 + e^- + \frac{1}{2}H^+ \rightarrow \frac{1}{2}C_6H_4ClNH_2 + \frac{1}{2}Cl^- \qquad \Delta G_a^{o\prime} = -28.76\ kJ/e^-eq$$

The $\Delta G_a^{o\prime}$ value was calculated from the standard Gibbs free energies of formation values reported in Susarla et al. (1997)[6] and Rittmann and McCarty (2001)[5],using the equation $\Delta G_a^{o\prime} = \sum \Delta G_{f(products)}^{o\prime} - \sum \Delta G_{f(reactants)}^{o\prime} = -28.76\ kJ/e^-eq.$

The $\Delta G_f^o$ values used to calculate the $\Delta G_a^{o\prime}$ are listed below.

| | $\Delta G_f^o$ | Reference |
|---|---|---|
| Aniline (aq) | -25.44 | Susarla et al. (1997)[6] |
| 2-CA (aq) | -69.51 | Susarla et al. (1997)[6] |
| 2,3-DCA (aq) | -105.38 | Susarla et al. (1997)[6] |
| $Cl^-$ (aq) | -133.26 | Rittmann and McCarty (2001)[5] |
| $H^+$ ($10^{-7}$ M) | -39.87 | Rittmann and McCarty (2001)[5] |

**iii.** The energy reaction ($R_{energy}$) is obtained using $R_e = R_a - R_d$ ($\Delta G_r^{o\prime} = \Delta G_a^{o\prime} - \Delta G_d^{o\prime}$ ):

$$R_{energy}: \frac{1}{2}C_6H_3Cl_2NH_2 + \frac{1}{2}H_2 \rightarrow \frac{1}{2}C_6H_4ClNH_2 + \frac{1}{2}H^+ + \frac{1}{2}Cl^- \qquad \Delta G_r^{o\prime} = -68.63\ kJ/e^-eq$$

Similarly, the $R_{energy}$ with 2-CA as the electron acceptor is also shown here:

$$\frac{1}{2}C_6H_4ClNH_2 + \frac{1}{2}H_2 \rightarrow \frac{1}{2}C_6H_5NH_2 + \frac{1}{2}H^+ + \frac{1}{2}Cl^- \qquad \Delta G_{r,2CA}^{o\prime} = -64.53\ kJ/e^-eq$$

**iv.** Assuming that ammonium serves as the nitrogen source for cell synthesis, the cell formation equation ($R_c$) is

$$R_c: \quad \frac{1}{5}CO_2 + \frac{1}{20}NH_4^+ + \frac{1}{20}HCO_3^- + H^+ + e^- \rightarrow \frac{1}{20}C_5H_7O_2N + \frac{9}{20}H_2O$$

Assuming ammonium is the nitrogen source, $\Delta G_{pc}^{o\prime} = \frac{113\ g\ cell}{1\ mol\ cell} \times \frac{1\ mol\ cell}{20\ e^{-}eq} \times 3.33 \frac{kJ}{g\ cell} =$ $18.81\ kJ/e^{-}eq$, where $\Delta G_{pc}^{o\prime}$ is the energy required for converting intermediate compound carbon to cellular carbon.

**v.** Assuming acetyl-CoA is the intermediate compound that is used for synthesizing macromolecules in cells, $\Delta G_{p}^{o\prime}$ is

$$\Delta G_{p}^{o\prime} = \Delta G_{CO2\rightarrow acet.CoA}^{o\prime} - \Delta G_{CO2\rightarrow CarbonSource}^{o\prime}$$

Given that the carbon source for the *Dehalobacter* is acetate,[7] the $\Delta G_{CO2\rightarrow CarbonSource}^{o\prime}$ can be calculated from

$$\frac{1}{8}CO_2 + \frac{1}{8}HCO_3^- + H^+ + e^- \rightarrow \frac{1}{8}CH_3COO^- + \frac{3}{8}H_20,\ \ \Delta G_{CO2\rightarrow CarbonSource}^{o\prime} = 27.40\ kJ/e^{-}eq$$

$$\Delta G_{p}^{o\prime} = \Delta G_{CO2\rightarrow acet.CoA}^{o\prime} - \Delta G_{CO2\rightarrow CarbonSource}^{o\prime} = 30.9 - 27.40 = 3.5\ kJ/e^{-}eq$$

In this example, the energy transfer efficiency is assumed to be 60%. The overall energy required for cell synthesis ($\Delta G_{S}^{o\prime}$) is

$$\Delta G_{S}^{o\prime} = \frac{\Delta G_{p}^{o\prime}}{\varepsilon^{n}} + \frac{\Delta G_{pc}^{o\prime}}{\varepsilon} = \frac{3.5}{0.6^{1}} + \frac{18.81}{0.6} = 37.18\ kJ/e^{-}eq$$

where an n value of 1 was employed due to a positive $\Delta G_{p}^{o\prime}$.

**vi.** The ratio of the energy required for synthesizing cells to the energy generated by dechlorination considering energy transfer losses, A, is

$$A = \frac{f_e}{f_s} = -\frac{\Delta G_{S}^{o\prime}}{\varepsilon \Delta G_{r}^{o\prime}} = \frac{-37.18\ kJ/e^{-}eq}{0.6 \times -68.63\ kJ/e^{-}eq} = 0.90$$

**vii.** The maximum value of growth yield in $\frac{eeq\ cell}{eeq\ donor}$, $f_s$, is

$$f_s = \frac{1}{1+A} = \frac{1}{1+0.90} = 0.53\ \frac{eeq\ cell}{eeq\ donor}$$

**viii.** The cell synthesis reaction ($R_{syn}$) is $R_{syn} = R_c - R_{carbon\ source}$

$$R_{syn}: \frac{1}{8}CH_3COO^- + \frac{3}{40}CO_2 + \frac{1}{20}NH_4^+ \rightarrow \frac{3}{40}HCO_3^- + \frac{1}{20}C_5H_7O_2N + \frac{3}{40}H_20$$

The energy reaction ($R_{energy}$) is $R_{energy} = R_a - R_d$

$$R_{\text{energy}}: \frac{1}{2}C_6H_3Cl_2NH_2 + \frac{1}{2}H_2 \rightarrow \frac{1}{2}C_6H_4ClNH_2 + \frac{1}{2}H^+ + \frac{1}{2}Cl^-$$

**ix.** The overall stoichiometric equation for *Dehalobacter* growing on 2,3-DCA is

$$R = R_{energy}f_e + R_{syn}f_s = R_{energy}(1 - f_s) + R_{syn}f_s = 0.47R_{energy} + 0.53R_{syn}$$

$$R: H_2 + C_6H_3Cl_2NH_2 + 0.28CH_3COO^- + 0.11NH_4^+ + 0.17CO_2$$
$$\rightarrow 0.11C_5H_7O_2N + C_6H_4ClNH_2 + Cl^- + 0.17HCO_3^- + 0.17H_20 + H^+$$

**i.** The growth yield, assuming an energy transfer efficiency of 60%, is

$$Y = 0.53\frac{eeq\ cell}{eeq\ donor} = 0.53\frac{eeq\ Dhb}{eeq\ H_2}$$

Since 1 eeq $H_2$ is equivalent to 1 eeq 2,3-DCA

$$Y = 0.53\frac{eeq\ Dhb}{eeq\ 2{,}3 - DCA} = 0.041\ g/g$$

Predicted yields at different ε for both dechlorination stages are summarized in Table S9.

**Section S10. Analysis of electron distribution and balances**

The electrons available from organic acids and alcohols were calculated as follows: 12 electron equivalents ($e^{-}$eq) for 1 mole of lactate, 12 $e^{-}$eq for 1 mole of ethanol, 14 $e^{-}$eq for 1 mole of propionate, 8 $e^{-}$eq for 1 mole of acetate, and 2 $e^{-}$eq for 1 mole of formate. Additionally, each mole of methane and hydrogen corresponds to 8 and 2 $e^{-}$eq, respectively. The $e^{-}$eq consumed for the dechlorination of 2,3-DCA were determined based on the measured consumption of 2,3-DCA, with 2 $e^{-}$eq required for transforming each mole of 2,3-DCA into CAs. Similarly, the electrons required for the dechlorination of 2-CA were calculated considering the measured aniline formation, which necessitates 2 $e^{-}$eq for the conversion of 2-CA to aniline per mole. The results of electron distribution for biotic replicates are presented in Figure S8. The electron balance (expressed in percentage) for each replicate was calculated as the sum of $e^{-}$eq for all components listed above on day 94 divided by the sum of $e^{-}$eq on day 0. The ethanol concentration on day 0 was calculated based on the amount of ethanol added into the bottles at time zero. It was assumed that ethanol was completely removed through fermentation after day zero. Headspace hydrogen gas on day 0 was estimated based on the hydrogen concentration measured in the glovebox on the day the bottles were set up and opened to the glovebox atmosphere (1.2% (v/v) hydrogen). It was also assumed that hydrogen concentration was low for the rest of time points due to consumption by *Dehalobacter* and methanogens. At some time points, samples for different analyses were not collected on the same day. To perform electron balances, data from nearby dates were used, as follows: Day 22: methane data measured on day 19 was used; Day 29: methane data measured on day 28 was used ;Day 36: methane data measured on day 39 was used; Day 50: methane data measured on day 49 was used; Day 94: 2,3-DCA and aniline concentration data measured on day 96 were used.

**Section S11. Stoichiometry of electron donor conversion during fermentation**

**The stoichiometry ratio of consumed lactate to formed propionate:**

The stoichiometry ratio was calculated based on the reduction of lactate and the formation of propionate between days 0 and 9. The average ratio across four biotic replicates was 2.94 ± 0.16:2 for consumed lactate to formed propionate, indicating fermentation of lactate to propionate and acetate. The stoichiometry is represented by the following reaction (Equation 3).[8,9]

$$\underset{\text{lactate}}{3\ C_3H_5O_3^-} \rightarrow \underset{\text{propionate}}{2\ C_3H_5O_2^-} + \underset{\text{acetate}}{C_2H_3O_2^-} + CO_2 + H_2O \qquad Eq.(3)$$

**Hydrogen-yielding and non-hydrogen-yielding acetogenesis:**

We calculated the ratio of ethanol (spiked at 3.3 mM at time zero) to measured acetate production, excluding the acetate formation attributed to lactate fermentation (where three moles of lactate produce one mole of acetate – see Eq. 3). Assuming all added ethanol was consumed, the ratio ethanol: acetate exclusively produced from ethanol was found to be 1:1.2 ± 0.4 (n=4). Equations 4 and 5 below show two common pathways of acetogenesis from ethanol. Acetogenesis from ethanol via the Wood-Ljungdahl pathway (ethanol:acetate = 1:1.5) is shown in Equation 4.[10]

$$\underset{\text{ethanol}}{2\ C_2H_6O} + 2\ CO_2 \rightarrow \underset{\text{acetate}}{3\ C_2H_3O_2^-} + 3H^+ \qquad Eq.(4)$$

Acetogenesis from ethanol to acetate and hydrogen (ethanol:acetate = 1:1) is shown in Equation 5.[8,10]

$$\underset{\text{ethanol}}{C_2H_6O} + H_2O \rightarrow \underset{\text{acetate}}{C_2H_3O_2^-} + H^+ + 2H_2 \qquad Eq.(5)$$

For biotic bottles I, II, and III, the ratio was calculated as 1:1.3 ± 0.4 using data from days 0 and 9, as no significant acetate production was observed after day 9. In contrast, for biotic bottle IV, the ratio was calculated as 1:0.9 using data from days 0 and 94, due to the slower acetogenesis observed in that bottle (Figure S6IV).

Therefore, our results aligned with the stoichiometry of fermentations of lactate and ethanol reported in the literature. We also measured formate, and its concentration was significantly lower than propionate and acetate (Figure S6). Formate formation was likely due to fermentative activities.

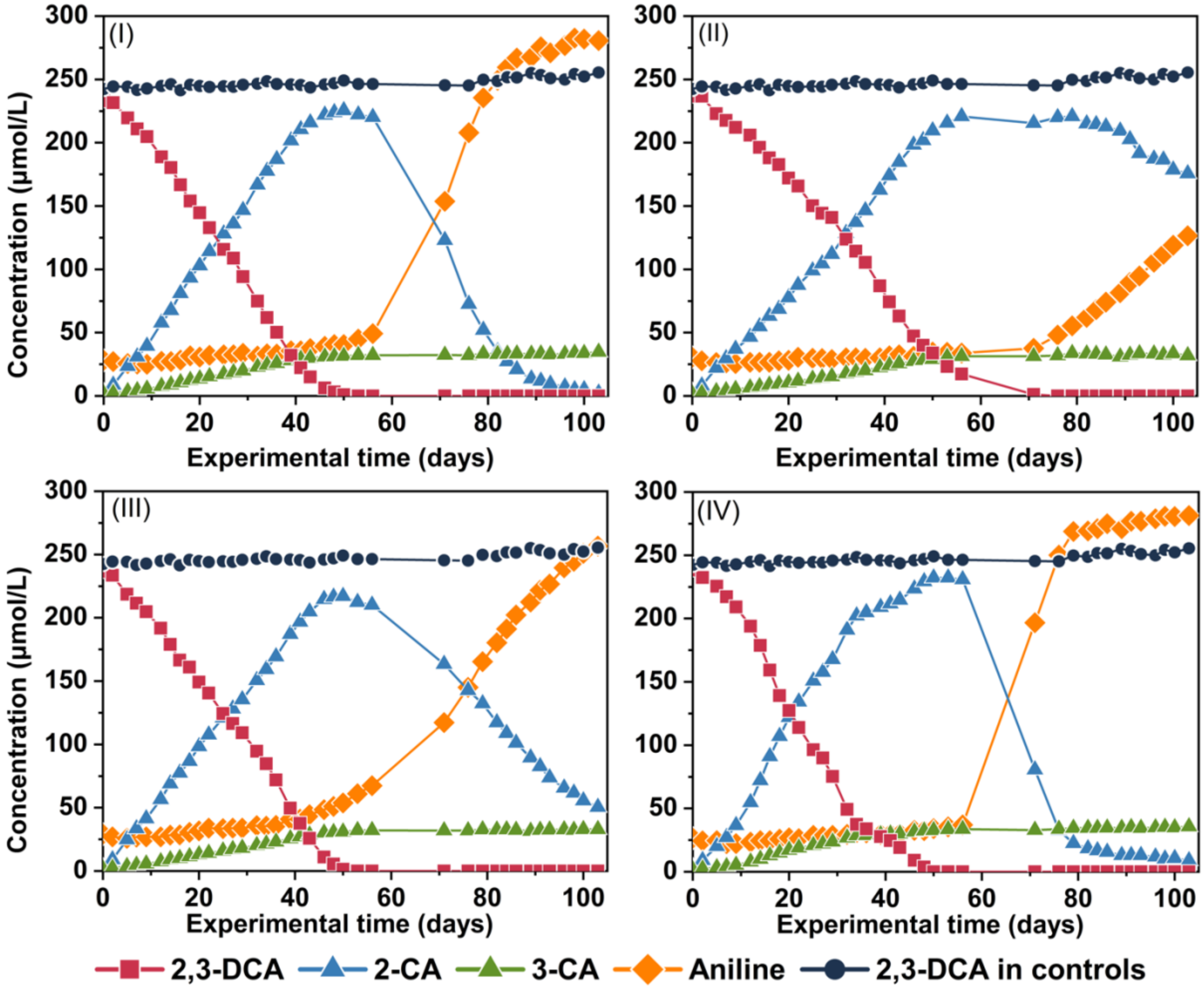


**Figure S5. Dechlorination of 2,3-DCA to aniline in four biotic replicates (bottles I to IV). The concentrations of chloroanilines and aniline are shown in the figure.**

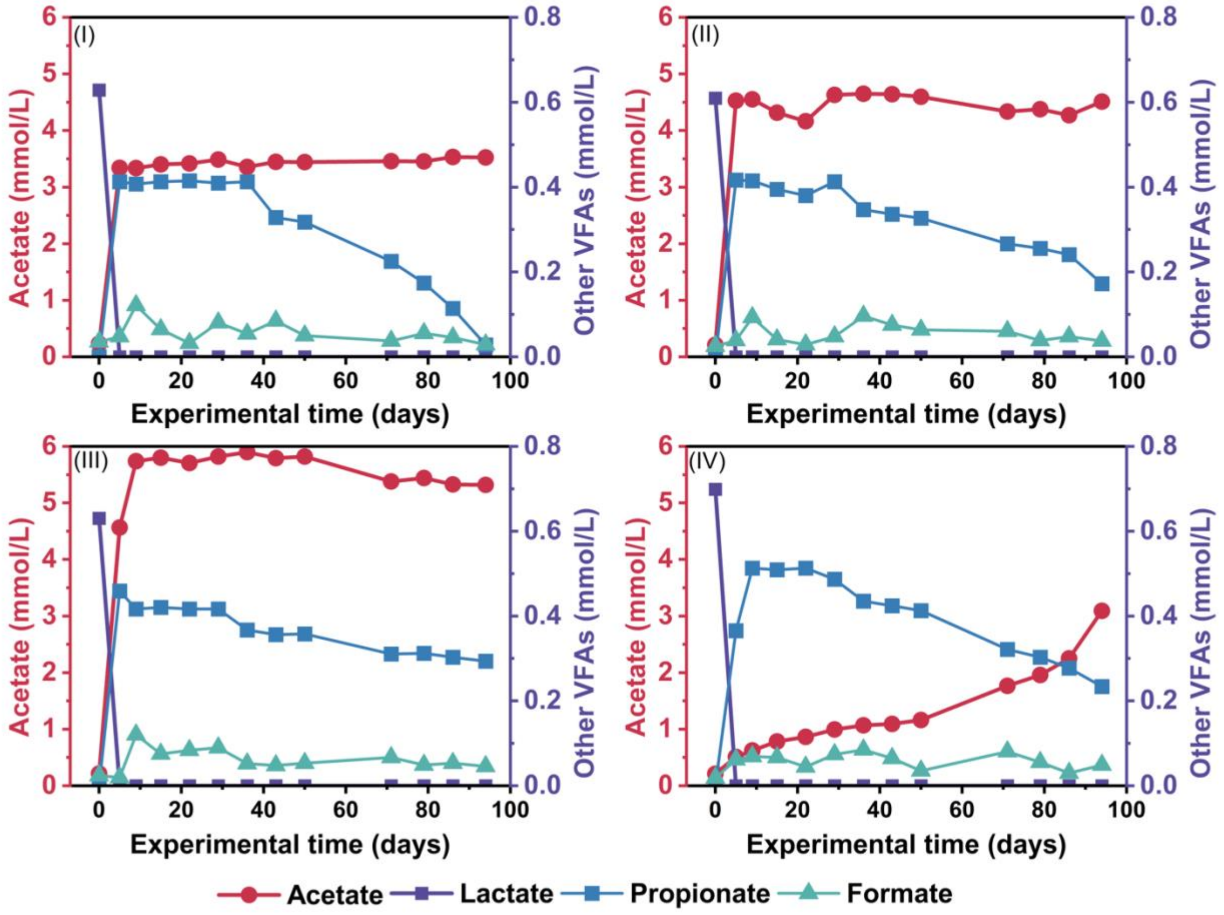


**Figure S6. Concentrations (mmol/L) of lactate (purple squares), acetate (red circles), propionate (blue squares), and formate (light teal triangles) in four biotic replicates (bottles I to IV). Butyrate and pyruvate were not detected. A consistent conversion of lactate to propionate at a 3:2 ratio is seen in this figure. Details are discussed in the main text and Section S11.**

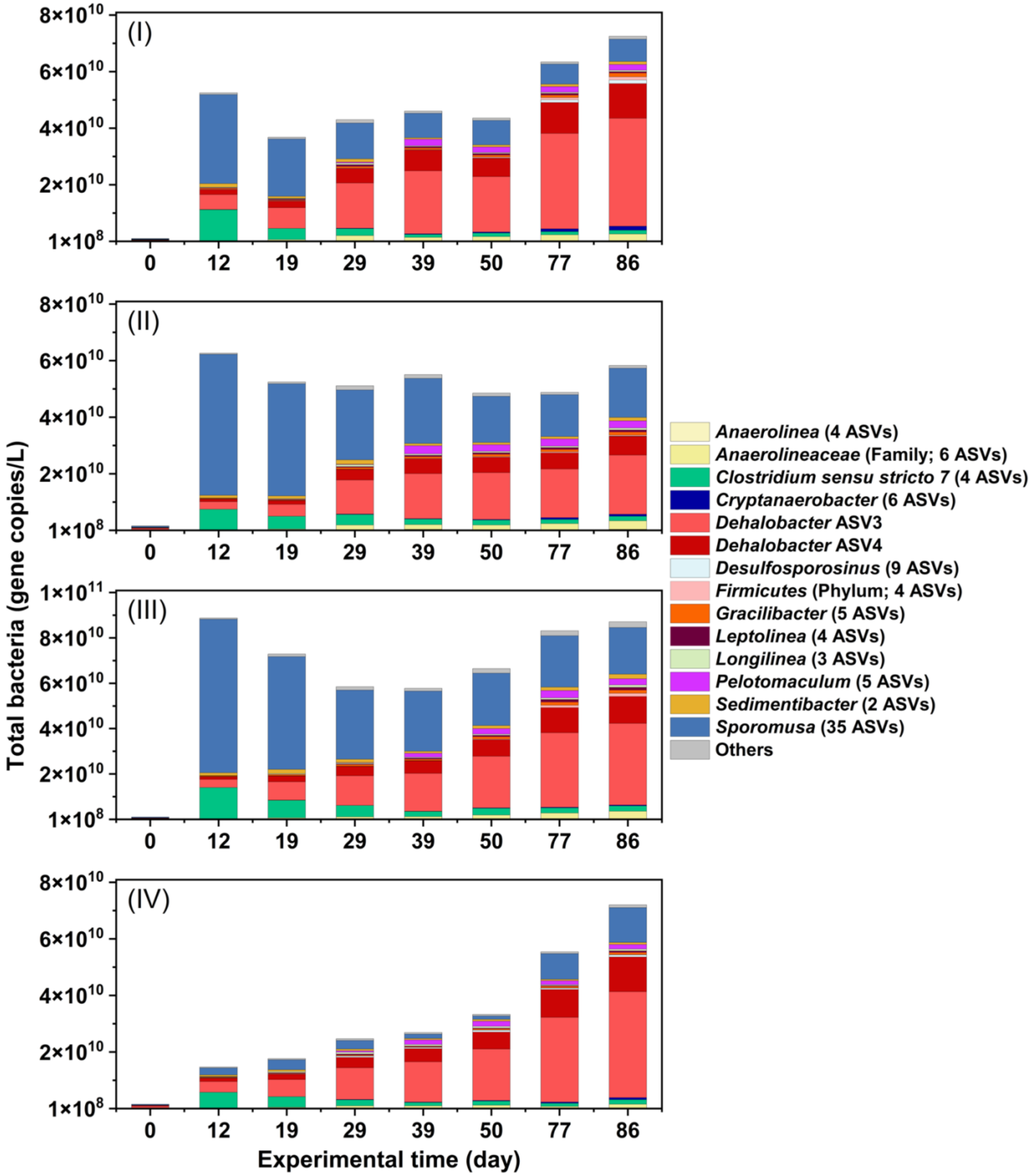


**Figure S7a. Relative abundance of bacteria at the genus level (determined from amplicon sequencing) combined with the absolute abundance of total bacteria**

**(quantified by qPCR targeting general bacteria, reported as 16S rRNA gene copies/L) for all biotic bottles (I to IV) during the course of the experiment.**

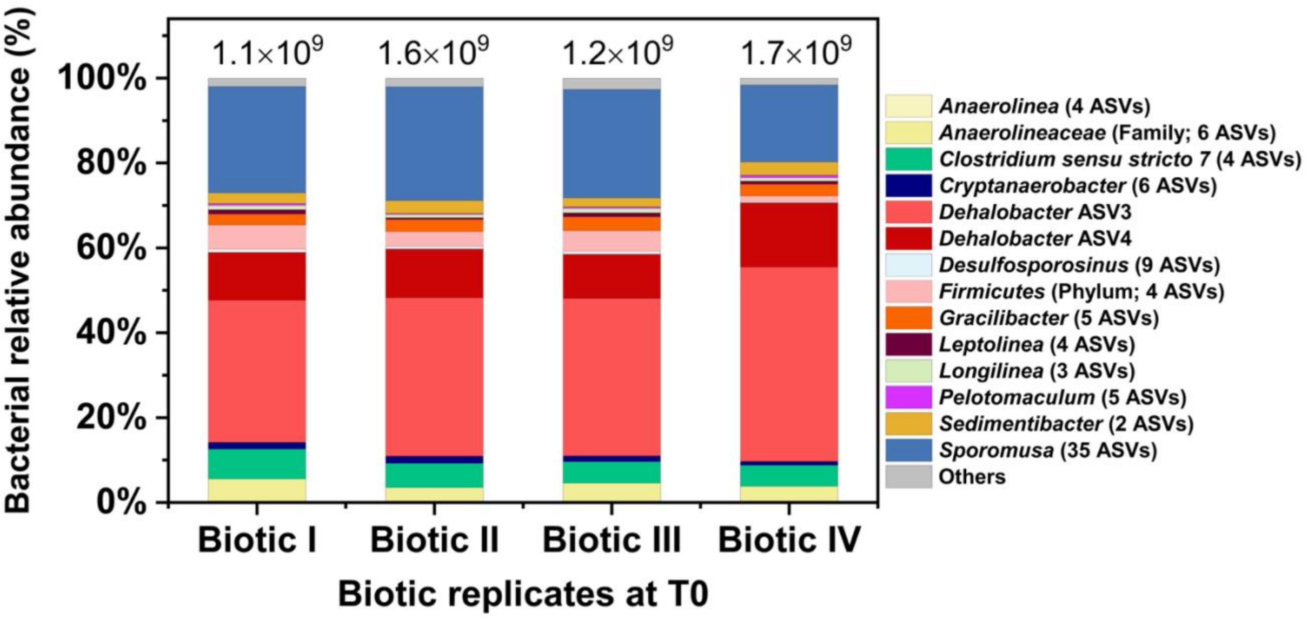


**Figure S7b. Initial bacterial compositions along with the absolute abundance of total bacteria (16S rRNA gene copies/L) for all four biotic replicates (bottles I to IV).**

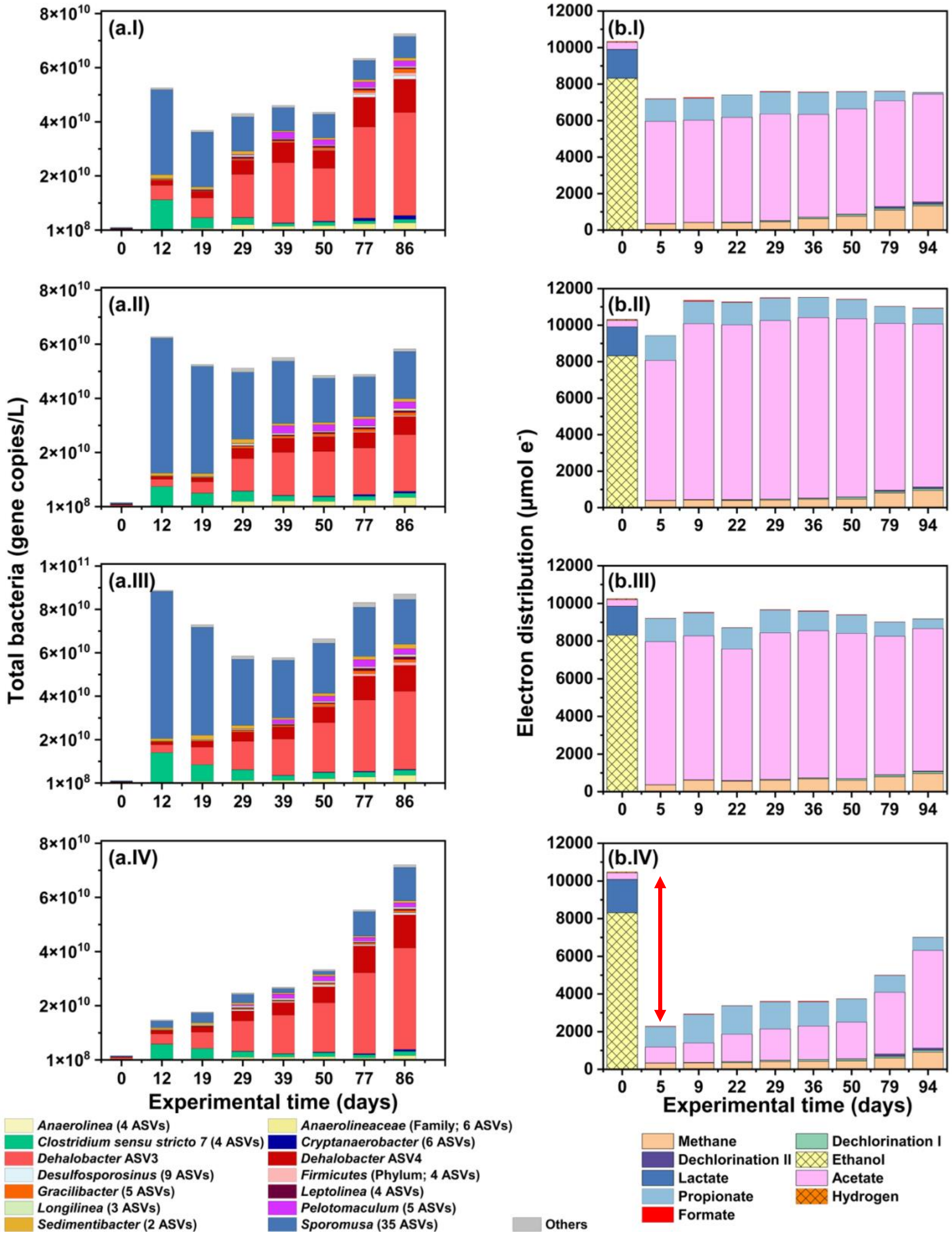

(a.I)
(a.II)
(a.III)
(a.IV)
(b.I)
(b.II)
(b.III)
(b.IV)
Total bacteria (gene copies/L)
Electron distribution (µmol e⁻)
Experimental time (days)
Anaerolinea (4 ASVs)
Anaerolineaceae (Family; 6 ASVs)
Clostridium sensu stricto 7 (4 ASVs)
Cryptanaerobacter (6 ASVs)
Dehalobacter ASV3
Dehalobacter ASV4
Desulfosporosinus (9 ASVs)
Firmicutes (Phylum; 4 ASVs)
Gracilibacter (5 ASVs)
Leptolinea (4 ASVs)
Longilinea (3 ASVs)
Pelotomaculum (5 ASVs)
Sedimentibacter (2 ASVs)
Sporomusa (35 ASVs)
Others
Methane
Dechlorination I
Dechlorination II
Ethanol
Lactate
Acetate
Propionate
Hydrogen
Formate

**Figure S8. Bacterial composition (panel a) alongside electron balances and distributions (panel b) for all four biotic replicates (I to IV) during reductive dechlorination of 2,3-DCA to aniline. Electrons stored in methane (including both headspace and dissolved forms), lactate, acetate, propionate, and formate, calculated based on the measured results (Section S11), are presented. Dechlorination I and Dechlorination II represent electron consumption for the reductive dechlorination of 2,3-DCA and 2-CA, respectively, as calculated from substrate depletion results. Electrons in ethanol at time zero were calculated based on the theoretical amount added to the system, while the initial headspace hydrogen was estimated using the hydrogen level measured in the glovebox atmosphere. Both ethanol and headspace hydrogen were assumed to be zero during the course of the experiment. The red double-ended arrow indicates the lack of electron balance in biotic replicate IV.**

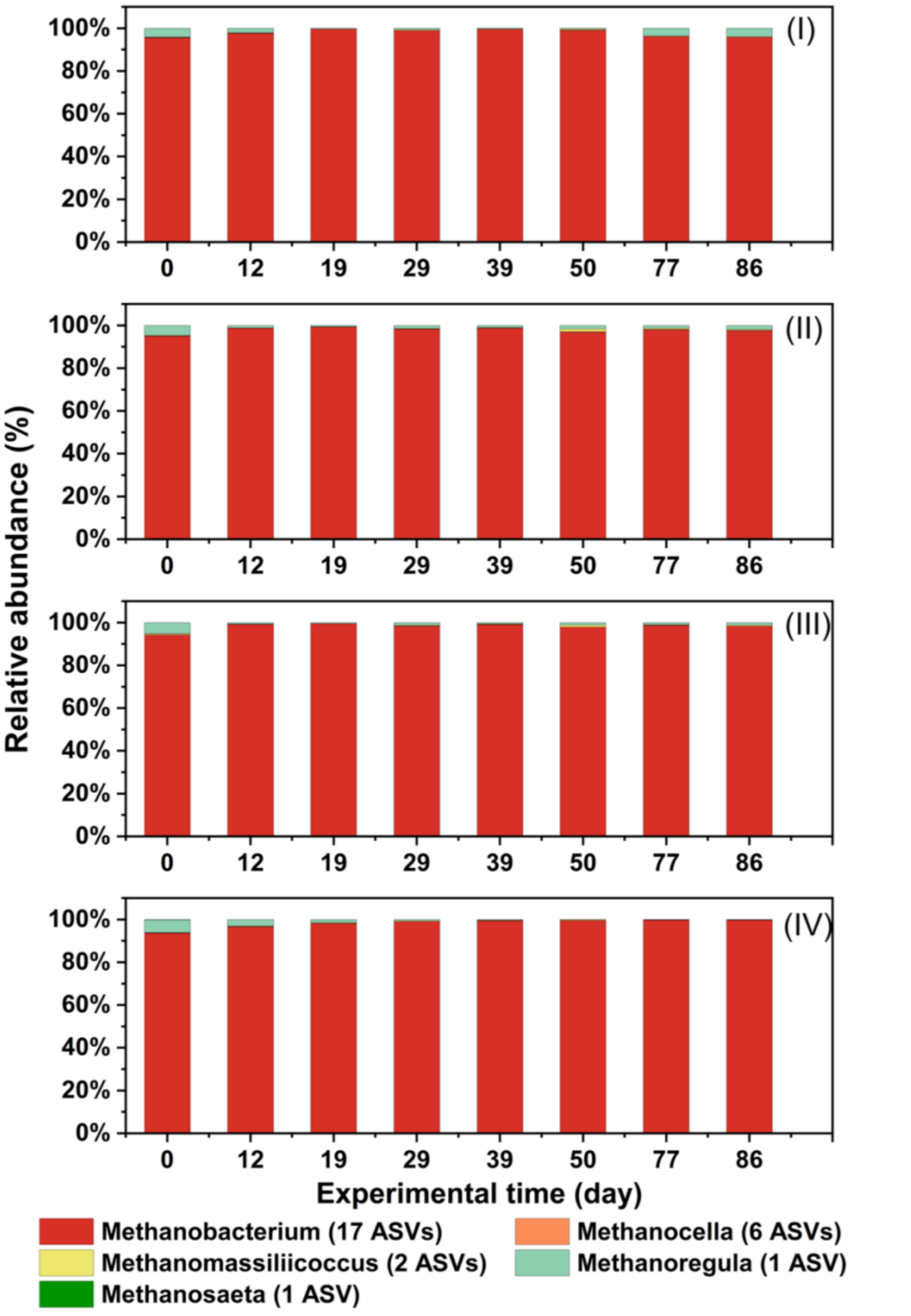


**Figure S9. Relative abundance of archaea at the genus level in biotic bottles (I to IV) during the course of the experiment.**

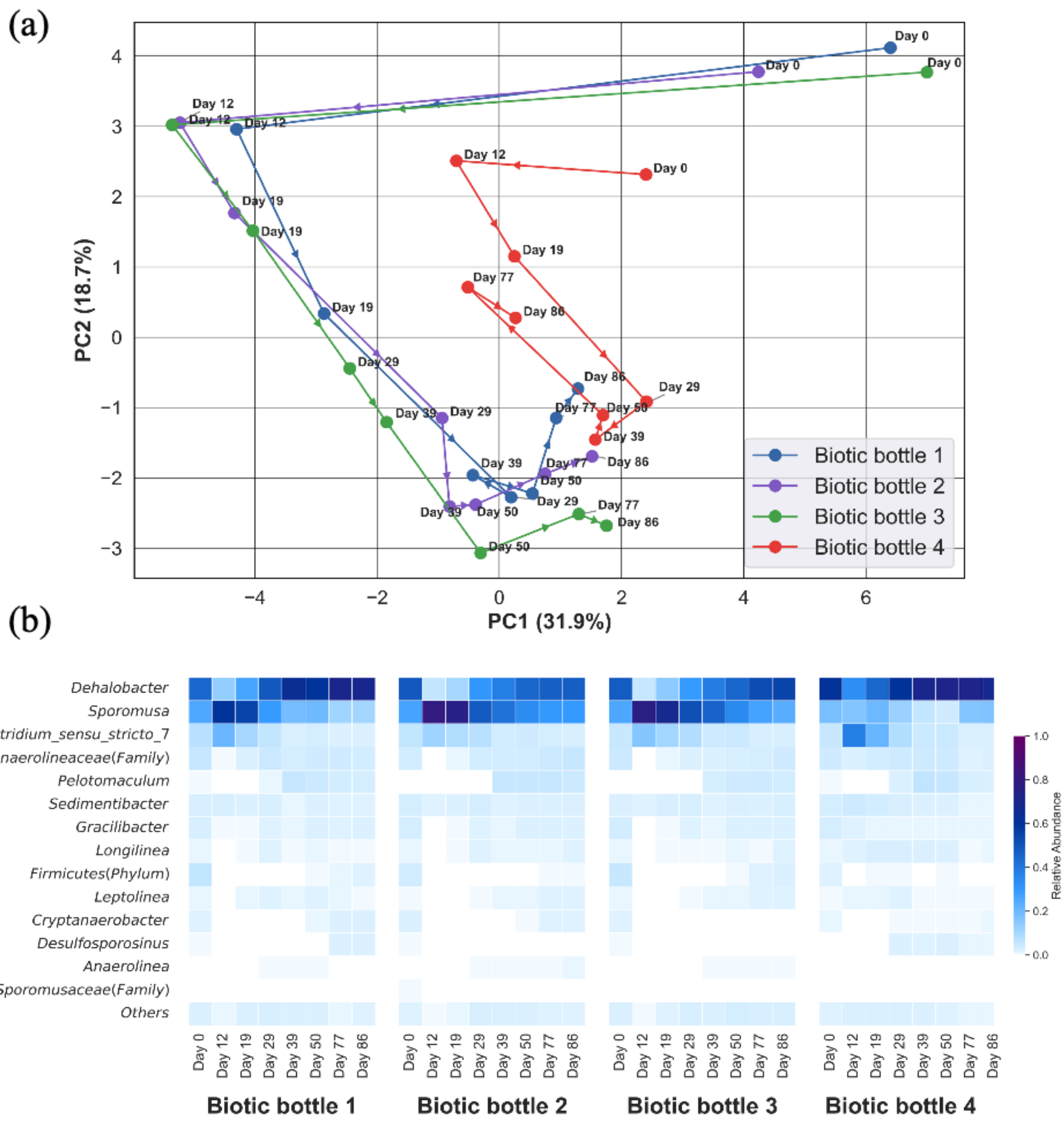


**Figure S10. (a) Principal component analysis (PCA) of bacterial community composition across biotic bottles during the incubation period. Each point represents a sample at a specific time point, colored by biotic bottle and labeled by day. Arrows indicate the temporal progression of microbial communities. (b) Heatmap of relative abundances of main bacterial genera over time. PCA was performed using the relative abundances of the 25 most abundant bacterial genera (accounting for >99.8% of the total), with the remaining genera (<0.2%) combined into a single 'Others' category. The analysis was**

**conducted using the PCA function from the sklearn.decomposition package in Python. The PCA results, combined with the genus-level heatmap, support the findings shown in the manuscript (Section 6.4). PCA reveals strong, time-driven, and reproducible bacterial succession. Biotic replicates I, II, and III exhibit similar temporal succession patterns, whereas replicate IV shows a slightly distinct community trajectory, likely due to limited growth of *Sporomusa* (panel b). Day 0 samples are clustered in the top-right quadrant, showing similar initial compositions (also shown in Panel b). Between Days 0 and 12, replicates I, II and III follow a similar trajectory driven by the dominance of fermentative bacteria (*Sporomusa* and *Clostridium sensu stricto 7*; panel b). From Day 12 onward, communities in replicates I, II and III shift toward the bottom-center region and converge across bottles at later time points, due to the increasing dominance of *Dehalobacter (panel b)*. This illustrates the functional adaptation of the communities driven by selective pressure from chlorinated aniline exposure.**

**Table S4. Sequence alignments for predominant *Dehalobacter* ASVs involved in chloroaniline dechlorination (ASVs 1 and 2 from the companion work[1]; ASVs 3 and 4 from this work).**

| % | ASV1 | ASV2 | ASV3 | ASV4 |
|---|---|---|---|---|
| ASV1 | / | 97.01 | 97.44 | 97.22 |
| ASV2 | 97.01 | / | 97.65 | 97.44 |
| ASV3 | 97.44 | 97.65 | / | 99.79 |
| ASV4 | 97.22 | 97.44 | 99.79 | / |

```
Consensus         GGGCCCGCACAAGCGGTGGAGCATGTGGTTTAATTCGACGCAACGCGAAGAACCTTACCA   60
Dehalobacter ASV1 GGGCCCGCACAAGCGGTGGAGCATGTGGTTTAATTCGACGCAACGCGAAGAACCTTACCA   60
Dehalobacter ASV2 GGGCCCGCACAAGCGGTGGAGCATGTGGTTTAATTCGACGCAACGCGAAGAACCTTACCA   60
Dehalobacter ASV3 GGGCCCGCACAAGCGGTGGAGCATGTGGTTTAATTCGACGCAACGCGAAGAACCTTACCA   60
Dehalobacter ASV4 GGGCCCGCACAAGCGGTGGAGCATGTGGTTTAATTCGACGCAACGCGAAGAACCTTACCA   60

Consensus         AGGCTTGACATCCATAGAATCCTGGAGAGATCCGGGAGTGCCCTTCGGGGARCTATGAGA  120
Dehalobacter ASV1 AGGCTTGACATCCAACTAATCCCGTAGAGATATGGGAGTGCCCTTCGGGGAAAGTTGAGA  120
Dehalobacter ASV2 AGGCTTGACATCTACAGAATCCTTAAGAGATTAGGGAGTGCCTTTCGGGGAACTGTAAGA  120
Dehalobacter ASV3 AGGCTTGACATCCATAGAATCCTGGAGAGATCCGGGAGTGCCCTTCGGGGAGCTATGAGA  120
Dehalobacter ASV4 AGGCTTGACATCCATAGAATCCTGGAGAGATCCGGGAGTGCCCTTCGGGGAGCTATGAGA  120

Consensus         CAGGTGGTGCATGGTTGTCGTCAGCTCGTGTCGTGAGATGTTGGGTTAAGTCCCGCAACG  180
Dehalobacter ASV1 CAGGTGGTGCATGGTTGTCGTCAGCTCGTGTCGTGAGATGTTGGGTTAAGTCCCGCAACG  180
Dehalobacter ASV2 CAGGTGGTGCATGGTTGTCGTCAGCTCGTGTCGTGAGATGTTGGGTTAAGTCCCGCAACG  180
Dehalobacter ASV3 CAGGTGGTGCATGGTTGTCGTCAGCTCGTGTCGTGAGATGTTGGGTTAAGTCCCGCAACG  180
Dehalobacter ASV4 CAGGTGGTGCATGGTTGTCGTCAGCTCGTGTCGTGAGATGTTGGGTTAAGTCCCGCAACG  180

Consensus         AGCGCAACCCCTATATTTAGTTGCTAACAGGTAAAGCTGAGAACTCTAGATAGACTGCCG  240
Dehalobacter ASV1 AGCGCAACCCCTATATTTAGTTGCTAACAGGTAAAGCTGAGAACTCTAGATAGACTGCCG  240
Dehalobacter ASV2 AGCGCAACCCCTATATTTAGTTGCTAACAGGTAAAGCTGAGAACTCTAGATAGACTGCCG  240
Dehalobacter ASV3 AGCGCAACCCCTATATTTAGTTGCTAACAGGTAAAGCTGAGAACTCTAGATAGACTGCCG  240
Dehalobacter ASV4 AGCGCAACCCCTATATTTAGTTGCTAACAGGTAAAGCTGAGAACTCTAGATAGACTGCCG  240

Consensus         GTGACAAACCGGAGGAAGGTGGGGATGACGTCAAATCATCATGCCCCTTATGTCTTGGGC  300
Dehalobacter ASV1 GTGACAAACCGGAGGAAGGTGGGGATGACGTCAAATCATCATGCCCCTTATGTCTTGGGC  300
Dehalobacter ASV2 GTGACAAACCGGAGGAAGGTGGGGATGACGTCAAATCATCATGCCCCTTATGTCTTGGGC  300
Dehalobacter ASV3 GTGACAAACCGGAGGAAGGTGGGGATGACGTCAAATCATCATGCCCCTTATGTCTTGGGC  300
Dehalobacter ASV4 GTGACAAACCGGAGGAAGGTGGGGACGACGTCAAATCATCATGCCCCTTATGTCTTGGGC  300

Consensus         TACACACGTGCTACAATGGACGGTACAGACGGAAGCGAAGCCGCGAGGTGRAGCAAATCC  360
Dehalobacter ASV1 TACACACGTGCTACAATGGACGGTACAGACGGAAGCGAAGCCGCGAGGTGAAGCAAATCC  360
Dehalobacter ASV2 TACACACGTGCTACAATGGACGGTACAGACGGAAGCGAAGCCGCGAGGTGAAGCAAATCC  360
Dehalobacter ASV3 TACACACGTGCTACAATGGACGGTACAGACGGAAGCGAAGCCGCGAGGTGGAGCAAATCC  360
Dehalobacter ASV4 TACACACGTGCTACAATGGACGGTACAGACGGAAGCGAAGCCGCGAGGTGGAGCAAATCC  360

Consensus         GAGAAAGCCGTTCTCAGTTCGGATTGCAGGCTGCAACTCGCCTGCATGAAGTCGGAATCG  420
Dehalobacter ASV1 GAGAAAGCCGTTCTCAGTTCGGATTGCAGGCTGCAACTCGCCTGCATGAAGTCGGAATCG  420
Dehalobacter ASV2 GAGAAAGCCGTTCTCAGTTCGGATTGCAGGCTGCAACTCGCCTGCATGAAGTCGGAATCG  420
Dehalobacter ASV3 GAGAAAGCCGTTCTCAGTTCGGATTGCAGGCTGCAACTCGCCTGCATGAAGTCGGAATCG  420
Dehalobacter ASV4 GAGAAAGCCGTTCTCAGTTCGGATTGCAGGCTGCAACTCGCCTGCATGAAGTCGGAATCG  420

Consensus         CTAGTAATCGCAGGTCAGCACACTGCGGTGAATACGTTCCCGGGCCTT              468
Dehalobacter ASV1 CTAGTAATCGCAGGTCAGCACACTGCGGTGAATACGTTCCCGGGCCTT              468
Dehalobacter ASV2 CTAGTAATCGCAGGTCAGCACACTGCGGTGAATACGTTCCCGGGCCTT              468
Dehalobacter ASV3 CTAGTAATCGCAGGTCAGCACACTGCGGTGAATACGTTCCCGGGCCTT              468
Dehalobacter ASV4 CTAGTAATCGCAGGTCAGCACACTGCGGTGAATACGTTCCCGGGCCTT              468
```

#### Table S5. Data for chlorinated amino aromatics analysis

| | Biotic bottle 1 (μmol/L) | | | | Biotic bottle 2 (μmol/L) | | | | Biotic bottle 3 (μmol/L) | | | | Biotic bottle 4 (μmol/L) | | | | Abotic controls MEAN (n=3) (μmol/L) | | | |
|---|---|---|---|---|---|---|---|---|---|---|---|---|---|---|---|---|---|---|---|---|
| Day | Aniline | 2-CA | 3-CA | 2,3-DCA | Aniline | 2-CA | 3-CA | 2,3-DCA | Aniline | 2-CA | 3-CA | 2,3-DCA | Aniline | 2-CA | 3-CA | 2,3-DCA | Aniline | 2-CA | 3-CA | 2,3-DCA |
| 0 | 28.1 | 2.3 | 1.5 | 233.1 | 30.6 | 2.7 | 1.6 | 236.8 | 29.1 | 2.4 | 1.6 | 236.6 | 26.7 | 3.0 | 1.6 | 238.4 | <LOQ | <LOQ | <LOQ | 242.4 |
| 2 | 27.2 | 8.6 | 2.2 | 231.5 | 27.8 | 8.1 | 2.1 | 236.7 | 26.9 | 9.0 | 2.3 | 233.5 | 24.4 | 9.2 | 2.4 | 232.4 | <LOQ | <LOQ | <LOQ | 244.5 |
| 5 | 25.6 | 23.4 | 3.6 | 219.5 | 26.3 | 22.1 | 3.5 | 223.0 | 25.8 | 24.8 | 3.8 | 218.6 | 24.6 | 20.0 | 3.3 | 225.4 | <LOQ | <LOQ | <LOQ | 244.2 |
| 7 | 25.3 | 30.9 | 4.4 | 210.9 | 26.0 | 29.5 | 4.3 | 217.5 | 26.7 | 33.5 | 4.7 | 211.5 | 22.2 | 27.1 | 4.1 | 217.1 | <LOQ | <LOQ | <LOQ | 241.6 |
| 9 | 25.1 | 39.4 | 5.4 | 204.8 | 25.6 | 37.0 | 5.2 | 212.1 | 27.1 | 41.3 | 5.7 | 204.7 | 22.3 | 36.7 | 5.3 | 209.0 | <LOQ | <LOQ | <LOQ | 242.8 |
| 12 | 27.4 | 57.7 | 7.7 | 193.9 | 26.6 | 46.6 | 6.5 | 206.0 | 27.1 | 56.5 | 7.5 | 191.8 | 23.4 | 54.7 | 7.9 | 188.8 | <LOQ | <LOQ | <LOQ | 244.9 |
| 14 | 27.2 | 67.6 | 9.0 | 180.3 | 26.6 | 54.7 | 7.5 | 196.3 | 28.4 | 68.9 | 9.0 | 179.0 | 24.6 | 72.1 | 10.1 | 178.8 | <LOQ | <LOQ | <LOQ | 246.2 |
| 16 | 28.4 | 81.0 | 10.7 | 166.8 | 26.7 | 63.1 | 8.6 | 188.1 | 28.5 | 77.5 | 10.1 | 166.6 | 24.7 | 91.1 | 12.7 | 159.3 | <LOQ | <LOQ | <LOQ | 241.5 |
| 18 | 31.8 | 93.2 | 12.4 | 153.9 | 27.9 | 68.7 | 9.4 | 182.7 | 30.6 | 86.8 | 11.4 | 161.1 | 25.8 | 107.0 | 15.0 | 139.1 | <LOQ | <LOQ | <LOQ | 245.8 |
| 20 | 30.8 | 102.9 | 13.7 | 144.7 | 28.2 | 77.7 | 10.5 | 172.1 | 31.7 | 98.3 | 12.9 | 149.1 | 26.7 | 122.0 | 17.1 | 127.3 | <LOQ | <LOQ | <LOQ | 244.9 |
| 22 | 32.1 | 114.2 | 15.4 | 132.9 | 30.6 | 87.5 | 11.9 | 165.8 | 33.9 | 107.6 | 14.2 | 140.4 | 26.3 | 134.1 | 18.9 | 113.9 | <LOQ | <LOQ | <LOQ | 244.1 |
| 25 | 32.5 | 128.1 | 17.2 | 115.6 | 29.8 | 98.9 | 13.5 | 150.2 | 33.5 | 121.1 | 16.1 | 124.4 | 28.7 | 150.7 | 21.0 | 96.4 | <LOQ | <LOQ | <LOQ | 244.6 |
| 27 | 32.5 | 136.0 | 18.5 | 108.8 | 29.6 | 104.5 | 14.3 | 144.3 | 33.9 | 128.2 | 17.1 | 116.6 | 27.8 | 157.7 | 22.1 | 89.8 | <LOQ | <LOQ | <LOQ | 244.3 |
| 29 | 33.8 | 146.6 | 20.0 | 94.1 | 28.9 | 111.9 | 15.4 | 141.0 | 33.9 | 135.4 | 18.1 | 109.2 | 28.8 | 167.7 | 23.6 | 75.4 | <LOQ | <LOQ | <LOQ | 245.7 |
| 32 | 31.6 | 166.8 | 22.8 | 75.0 | 30.3 | 128.6 | 17.6 | 123.7 | 35.8 | 150.6 | 20.3 | 94.7 | 29.5 | 190.9 | 26.8 | 49.3 | <LOQ | <LOQ | <LOQ | 246.6 |
| 34 | 33.3 | 177.5 | 24.4 | 61.9 | 30.2 | 137.3 | 18.9 | 114.4 | 35.5 | 159.2 | 21.5 | 84.8 | 30.9 | 202.2 | 28.6 | 37.5 | <LOQ | <LOQ | <LOQ | 248.6 |
| 36 | 33.2 | 186.8 | 25.8 | 50.4 | 29.7 | 146.5 | 20.2 | 105.4 | 36.1 | 169.4 | 23.0 | 71.9 | 30.8 | 204.7 | 28.8 | 33.9 | <LOQ | <LOQ | <LOQ | 246.3 |
| 39 | 35.7 | 201.7 | 28.0 | 32.0 | 30.2 | 162.6 | 22.4 | 86.9 | 40.9 | 187.1 | 25.7 | 49.7 | 29.5 | 208.9 | 29.4 | 27.9 | <LOQ | <LOQ | <LOQ | 246.0 |
| 41 | 35.7 | 210.5 | 29.2 | 22.2 | 31.8 | 174.1 | 24.1 | 74.4 | 40.9 | 196.8 | 27.2 | 37.6 | 29.9 | 211.7 | 29.9 | 24.6 | <LOQ | <LOQ | <LOQ | 245.7 |
| 43 | 37.5 | 215.9 | 30.0 | 14.9 | 32.5 | 184.7 | 25.6 | 63.2 | 44.5 | 204.8 | 28.5 | 25.9 | 29.9 | 214.6 | 30.0 | 19.2 | <LOQ | <LOQ | <LOQ | 243.6 |
| 46 | 39.3 | 221.9 | 30.9 | 6.2 | 31.3 | 198.3 | 27.5 | 47.2 | 48.4 | 214.6 | 30.2 | 11.0 | 32.7 | 223.6 | 31.6 | 9.2 | <LOQ | <LOQ | <LOQ | 246.0 |
| 48 | 40.5 | 224.0 | 31.2 | 3.2 | 32.2 | 201.8 | 28.1 | 40.0 | 51.0 | 216.5 | 30.6 | 5.4 | 32.5 | 229.2 | 32.3 | 3.8 | <LOQ | <LOQ | <LOQ | 246.9 |
| 50 | 40.6 | 225.4 | 31.5 | 1.0 | 34.6 | 209.5 | 28.9 | 33.9 | 53.9 | 217.1 | 31.0 | 1.4 | 34.2 | 232.1 | 32.7 | <LOQ | <LOQ | <LOQ | <LOQ | 249.1 |
| 53 | 45.0 | 222.1 | 32.1 | <LOQ | 34.5 | 215.7 | 30.7 | 23.3 | 61.2 | 212.3 | 32.1 | <LOQ | 35.5 | 232.2 | 33.5 | <LOQ | <LOQ | <LOQ | <LOQ | 246.3 |
| 56 | 49.4 | 220.1 | 32.2 | <LOQ | 33.9 | 220.8 | 31.3 | 17.3 | 67.4 | 210.2 | 32.2 | <LOQ | 37.0 | 230.7 | 33.7 | <LOQ | <LOQ | <LOQ | <LOQ | 246.4 |
| 71 | 153.7 | 123.0 | 32.4 | <LOQ | 37.6 | 215.5 | 31.4 | 1.1 | 117.3 | 163.3 | 32.0 | <LOQ | 196.8 | 80.7 | 33.0 | <LOQ | <LOQ | <LOQ | <LOQ | 245.5 |
| 76 | 207.9 | 72.5 | 32.1 | <LOQ | 48.1 | 220.0 | 32.0 | <LOQ | 145.0 | 142.7 | 31.9 | <LOQ | 250.2 | 32.6 | 33.7 | <LOQ | <LOQ | <LOQ | <LOQ | 245.1 |
| 79 | 235.6 | 52.0 | 33.2 | <LOQ | 55.5 | 220.7 | 33.7 | <LOQ | 165.5 | 132.3 | 32.7 | <LOQ | 268.8 | 22.5 | 34.5 | <LOQ | <LOQ | <LOQ | <LOQ | 249.7 |
| 82 | 249.3 | 35.0 | 33.0 | <LOQ | 62.1 | 215.1 | 33.3 | <LOQ | 180.3 | 117.3 | 32.5 | <LOQ | 269.3 | 18.4 | 34.5 | <LOQ | <LOQ | <LOQ | <LOQ | 248.8 |
| 84 | 259.6 | 27.2 | 33.0 | <LOQ | 68.3 | 214.4 | 33.0 | <LOQ | 191.2 | 108.9 | 32.7 | <LOQ | 271.1 | 16.8 | 34.5 | <LOQ | <LOQ | <LOQ | <LOQ | 251.8 |
| 86 | 266.9 | 20.9 | 33.3 | <LOQ | 74.2 | 212.8 | 32.5 | <LOQ | 202.2 | 101.2 | 32.3 | <LOQ | 274.9 | 15.5 | 34.8 | <LOQ | <LOQ | <LOQ | <LOQ | 251.5 |
| 89 | 267.6 | 13.6 | 33.2 | <LOQ | 81.7 | 209.4 | 30.9 | <LOQ | 212.5 | 89.6 | 31.4 | <LOQ | 271.0 | 12.9 | 34.4 | <LOQ | <LOQ | <LOQ | <LOQ | 255.1 |
| 91 | 275.8 | 11.9 | 33.3 | <LOQ | 89.6 | 202.7 | 32.2 | <LOQ | 221.7 | 82.7 | 32.2 | <LOQ | 277.0 | 13.0 | 35.0 | <LOQ | <LOQ | <LOQ | <LOQ | 253.5 |
| 93 | 270.9 | 9.5 | 32.7 | <LOQ | 94.9 | 191.6 | 33.4 | <LOQ | 226.6 | 73.7 | 32.5 | <LOQ | 277.0 | 12.9 | 34.6 | <LOQ | <LOQ | <LOQ | <LOQ | 251.0 |
| 96 | 276.8 | 7.2 | 33.2 | <LOQ | 105.7 | 187.2 | 33.3 | <LOQ | 239.5 | 65.6 | 32.8 | <LOQ | 278.9 | 11.7 | 34.9 | <LOQ | <LOQ | <LOQ | <LOQ | 249.9 |
| 98 | 282.7 | 5.2 | 33.9 | <LOQ | 111.9 | 186.5 | 32.2 | <LOQ | 244.6 | 61.8 | 32.4 | <LOQ | 280.8 | 10.3 | 35.3 | <LOQ | <LOQ | <LOQ | <LOQ | 254.3 |
| 100 | 281.8 | 5.2 | 33.6 | <LOQ | 119.0 | 178.6 | 33.4 | <LOQ | 251.0 | 55.6 | 32.8 | <LOQ | 280.7 | 10.7 | 35.0 | <LOQ | <LOQ | <LOQ | <LOQ | 252.3 |
| 103 | 280.7 | 2.3 | 34.8 | <LOQ | 126.5 | 175.6 | 31.9 | <LOQ | 256.7 | 50.6 | 32.9 | <LOQ | 281.5 | 8.8 | 35.6 | <LOQ | <LOQ | <LOQ | <LOQ | 255.6 |

<LOQ: below the limit of quantificaiton.

LOQ (μmol/L) for chlorinated anilines and aniline: 2,3-DCA: 0.7; 2-CA: 0.8; 3-CA: 0.8; 4-CA: 3.9; aniline: 1.5.

**Table S6. Data for analysis of volatile fatty acids and inorganic anions (mmol/L)**

| Lactate (mmol/L) | | | | | | | | | | | | | |
|---|---|---|---|---|---|---|---|---|---|---|---|---|---|
| Day | 0 | 5 | 9 | 15 | 22 | 29 | 36 | 43 | 50 | 71 | 79 | 86 | 94 |
| Biotic bottle 1 | 0.63 | <LOQ | <LOQ | <LOQ | <LOQ | <LOQ | <LOQ | <LOQ | <LOQ | <LOQ | <LOQ | <LOQ | <LOQ |
| Biotic bottle 2 | 0.61 | <LOQ | <LOQ | <LOQ | <LOQ | <LOQ | <LOQ | <LOQ | <LOQ | <LOQ | <LOQ | <LOQ | <LOQ |
| Biotic bottle 3 | 0.63 | <LOQ | <LOQ | <LOQ | <LOQ | <LOQ | <LOQ | <LOQ | <LOQ | <LOQ | <LOQ | <LOQ | <LOQ |
| Biotic bottle 4 | 0.70 | <LOQ | <LOQ | <LOQ | <LOQ | <LOQ | <LOQ | <LOQ | <LOQ | <LOQ | <LOQ | <LOQ | <LOQ |
| Abiotic control MEAN (n=3) | <LOQ | <LOQ | <LOQ | <LOQ | <LOQ | <LOQ | <LOQ | <LOQ | <LOQ | <LOQ | <LOQ | <LOQ | <LOQ |
| Acetate (mmol/L) | | | | | | | | | | | | | |
| Day | 0 | 5 | 9 | 15 | 22 | 29 | 36 | 43 | 50 | 71 | 79 | 86 | 94 |
| Biotic bottle 1 | 0.23 | 3.34 | 3.34 | 3.40 | 3.42 | 3.49 | 3.36 | 3.45 | 3.44 | 3.46 | 3.45 | 3.53 | 3.52 |
| Biotic bottle 2 | 0.21 | 4.53 | 4.55 | 4.32 | 4.16 | 4.63 | 4.65 | 4.64 | 4.60 | 4.33 | 4.38 | 4.27 | 4.51 |
| Biotic bottle 3 | 0.22 | 4.57 | 5.74 | 5.80 | 5.71 | 5.82 | 5.90 | 5.79 | 5.82 | 5.38 | 5.44 | 5.33 | 5.32 |
| Biotic bottle 4 | 0.21 | 0.51 | 0.62 | 0.79 | 0.87 | 1.00 | 1.07 | 1.09 | 1.16 | 1.77 | 1.96 | 2.25 | 3.09 |
| Abiotic control MEAN (n=3) | <LOQ | <LOQ | <LOQ | <LOQ | <LOQ | <LOQ | <LOQ | <LOQ | <LOQ | <LOQ | <LOQ | <LOQ | <LOQ |
| Propionate (mmol/L) | | | | | | | | | | | | | |
| Day | 0 | 5 | 9 | 15 | 22 | 29 | 36 | 43 | 50 | 71 | 79 | 86 | 94 |
| Biotic bottle 1 | <LOQ | 0.41 | 0.41 | 0.41 | 0.41 | 0.41 | 0.41 | 0.33 | 0.32 | 0.23 | 0.17 | 0.11 | 0.03 |
| Biotic bottle 2 | <LOQ | 0.42 | 0.41 | 0.39 | 0.38 | 0.41 | 0.35 | 0.34 | 0.33 | 0.27 | 0.26 | 0.24 | 0.17 |
| Biotic bottle 3 | <LOQ | 0.46 | 0.42 | 0.42 | 0.42 | 0.42 | 0.37 | 0.36 | 0.36 | 0.31 | 0.31 | 0.30 | 0.29 |
| Biotic bottle 4 | <LOQ | 0.37 | 0.51 | 0.51 | 0.51 | 0.49 | 0.44 | 0.42 | 0.41 | 0.32 | 0.30 | 0.28 | 0.23 |
| Control MEAN (n=3) | <LOQ | <LOQ | <LOQ | <LOQ | <LOQ | <LOQ | <LOQ | <LOQ | <LOQ | <LOQ | <LOQ | <LOQ | <LOQ |
| Formate (mmol/L) | | | | | | | | | | | | | |
| Day | 0 | 5 | 9 | 15 | 22 | 29 | 36 | 43 | 50 | 71 | 79 | 86 | 94 |
| Biotic bottle 1 | 0.04 | 0.05 | 0.12 | 0.06 | 0.03 | 0.08 | 0.05 | 0.08 | 0.05 | 0.04 | 0.06 | 0.05 | 0.03 |
| Biotic bottle 2 | 0.03 | 0.04 | 0.09 | 0.04 | 0.03 | 0.05 | 0.10 | 0.07 | 0.06 | 0.06 | 0.04 | 0.05 | 0.04 |
| Biotic bottle 3 | 0.02 | 0.02 | 0.12 | 0.07 | 0.08 | 0.09 | 0.05 | 0.05 | 0.05 | 0.07 | 0.05 | 0.05 | 0.05 |
| Biotic bottle 4 | 0.02 | 0.06 | 0.07 | 0.07 | 0.04 | 0.07 | 0.08 | 0.06 | 0.04 | 0.08 | 0.06 | 0.03 | 0.05 |
| Control MEAN (n=3) | <LOQ | <LOQ | <LOQ | <LOQ | <LOQ | <LOQ | <LOQ | <LOQ | <LOQ | <LOQ | <LOQ | <LOQ | <LOQ |
| Butyrate (mmol/L) | | | | | | | | | | | | | |
| Day | 0 | 5 | 9 | 15 | 22 | 29 | 36 | 43 | 50 | 71 | 79 | 86 | 94 |
| Biotic bottle 1 | <LOQ | <LOQ | <LOQ | <LOQ | <LOQ | <LOQ | <LOQ | <LOQ | <LOQ | <LOQ | <LOQ | <LOQ | <LOQ |
| Biotic bottle 2 | <LOQ | <LOQ | <LOQ | <LOQ | <LOQ | <LOQ | <LOQ | <LOQ | <LOQ | <LOQ | <LOQ | <LOQ | <LOQ |
| Biotic bottle 3 | <LOQ | <LOQ | <LOQ | <LOQ | <LOQ | <LOQ | <LOQ | <LOQ | <LOQ | <LOQ | <LOQ | <LOQ | <LOQ |
| Biotic bottle 4 | <LOQ | <LOQ | <LOQ | <LOQ | <LOQ | <LOQ | <LOQ | <LOQ | <LOQ | <LOQ | <LOQ | <LOQ | <LOQ |
| Control MEAN (n=3) | <LOQ | <LOQ | <LOQ | <LOQ | <LOQ | <LOQ | <LOQ | <LOQ | <LOQ | <LOQ | <LOQ | <LOQ | <LOQ |
| Pyruvate (mmol/L) | | | | | | | | | | | | | |
| Day | 0 | 5 | 9 | 15 | 22 | 29 | 36 | 43 | 50 | 71 | 79 | 86 | 94 |
| Biotic bottle 1 | <LOQ | <LOQ | <LOQ | <LOQ | <LOQ | <LOQ | <LOQ | <LOQ | <LOQ | <LOQ | <LOQ | <LOQ | <LOQ |
| Biotic bottle 2 | <LOQ | <LOQ | <LOQ | <LOQ | <LOQ | <LOQ | <LOQ | <LOQ | <LOQ | <LOQ | <LOQ | <LOQ | <LOQ |
| Biotic bottle 3 | <LOQ | <LOQ | <LOQ | <LOQ | <LOQ | <LOQ | <LOQ | <LOQ | <LOQ | <LOQ | <LOQ | <LOQ | <LOQ |
| Biotic bottle 4 | <LOQ | <LOQ | <LOQ | <LOQ | <LOQ | <LOQ | <LOQ | <LOQ | <LOQ | <LOQ | <LOQ | <LOQ | <LOQ |
| Control MEAN (n=3) | <LOQ | <LOQ | <LOQ | <LOQ | <LOQ | <LOQ | <LOQ | <LOQ | <LOQ | <LOQ | <LOQ | <LOQ | <LOQ |
| [illegible] | | | | | | | | | | | | | |

| Chloride (mmol/L) | | | | | | | | | | | | | |
|---|---|---|---|---|---|---|---|---|---|---|---|---|---|
| Day | 0 | 5 | 9 | 15 | 22 | 29 | 36 | 43 | 50 | 71 | 79 | 86 | 94 |
| Biotic bottle 1 | 12.44 | 12.34 | 12.22 | 12.42 | 12.53 | 12.68 | 12.29 | 12.45 | 12.44 | 12.31 | 12.77 | 12.94 | 12.93 |
| Biotic bottle 2 | 11.98 | 11.66 | 12.32 | 11.77 | 11.42 | 12.56 | 12.52 | 12.50 | 12.44 | 12.58 | 12.76 | 12.97 | 12.83 |
| Biotic bottle 3 | 12.15 | 12.41 | 12.40 | 12.55 | 12.41 | 12.60 | 12.72 | 12.61 | 12.66 | 12.74 | 12.97 | 12.88 | 13.08 |
| Biotic bottle 4 | 12.25 | 12.15 | 12.46 | 12.46 | 12.70 | 12.78 | 12.80 | 12.60 | 12.56 | 12.74 | 12.81 | 12.95 | 13.22 |
| Abiotic control MEAN (n=3) | 12.60 | 12.46 | 12.53 | 12.38 | 12.52 | 12.53 | 12.44 | 12.43 | 12.44 | 12.41 | 12.61 | 12.34 | 12.37 |
| Nitrite (mmol/L) | | | | | | | | | | | | | |
| Day | 0 | 5 | 9 | 15 | 22 | 29 | 36 | 43 | 50 | 71 | 79 | 86 | 94 |
| Biotic bottle 1 | <LOQ | <LOQ | <LOQ | <LOQ | <LOQ | <LOQ | <LOQ | <LOQ | <LOQ | <LOQ | <LOQ | <LOQ | <LOQ |
| Biotic bottle 2 | <LOQ | <LOQ | <LOQ | <LOQ | <LOQ | <LOQ | <LOQ | <LOQ | <LOQ | <LOQ | <LOQ | <LOQ | <LOQ |
| Biotic bottle 3 | <LOQ | <LOQ | <LOQ | <LOQ | <LOQ | <LOQ | <LOQ | <LOQ | <LOQ | <LOQ | <LOQ | <LOQ | <LOQ |
| Biotic bottle 4 | <LOQ | <LOQ | <LOQ | <LOQ | <LOQ | <LOQ | <LOQ | <LOQ | <LOQ | <LOQ | <LOQ | <LOQ | <LOQ |
| Abiotic control MEAN (n=3) | <LOQ | <LOQ | <LOQ | <LOQ | <LOQ | <LOQ | <LOQ | <LOQ | <LOQ | <LOQ | <LOQ | <LOQ | <LOQ |
| Nitrate (mmol/L) | | | | | | | | | | | | | |
| Day | 0 | 5 | 9 | 15 | 22 | 29 | 36 | 43 | 50 | 71 | 79 | 86 | 94 |
| Biotic bottle 1 | <LOQ | <LOQ | <LOQ | <LOQ | <LOQ | <LOQ | <LOQ | <LOQ | <LOQ | <LOQ | <LOQ | <LOQ | <LOQ |
| Biotic bottle 2 | <LOQ | <LOQ | <LOQ | <LOQ | <LOQ | <LOQ | <LOQ | <LOQ | <LOQ | <LOQ | <LOQ | <LOQ | <LOQ |
| Biotic bottle 3 | <LOQ | <LOQ | <LOQ | <LOQ | <LOQ | <LOQ | <LOQ | <LOQ | <LOQ | <LOQ | <LOQ | <LOQ | <LOQ |
| Biotic bottle 4 | <LOQ | <LOQ | <LOQ | <LOQ | <LOQ | <LOQ | <LOQ | <LOQ | <LOQ | <LOQ | <LOQ | <LOQ | <LOQ |
| Abiotic control MEAN (n=3) | <LOQ | <LOQ | <LOQ | <LOQ | <LOQ | <LOQ | <LOQ | <LOQ | <LOQ | <LOQ | <LOQ | <LOQ | <LOQ |
| Sulfate (mmol/L) | | | | | | | | | | | | | |
| Day | 0 | 5 | 9 | 15 | 22 | 29 | 36 | 43 | 50 | 71 | 79 | 86 | 94 |
| Biotic bottle 1 | 0.06 | 0.05 | 0.06 | 0.06 | 0.07 | 0.06 | 0.03 | 0.03 | 0.03 | 0.04 | 0.03 | 0.03 | 0.03 |
| Biotic bottle 2 | 0.06 | 0.06 | 0.06 | 0.06 | 0.07 | 0.07 | 0.03 | 0.03 | 0.03 | 0.04 | 0.03 | 0.03 | 0.02 |
| Biotic bottle 3 | 0.06 | 0.05 | 0.06 | 0.06 | 0.07 | 0.07 | 0.04 | 0.04 | 0.03 | 0.03 | 0.03 | 0.02 | 0.03 |
| Biotic bottle 4 | 0.06 | 0.05 | 0.07 | 0.06 | 0.07 | 0.05 | 0.03 | 0.04 | 0.03 | 0.03 | 0.03 | 0.03 | 0.03 |
| Abiotic control MEAN (n=3) | 0.06 | 0.05 | 0.05 | 0.06 | 0.06 | 0.05 | 0.05 | 0.05 | 0.05 | 0.05 | 0.05 | 0.05 | 0.05 |
| Phosphate (mmol/L) | | | | | | | | | | | | | |
| Day | 0 | 5 | 9 | 15 | 22 | 29 | 36 | 43 | 50 | 71 | 79 | 86 | 94 |
| Biotic bottle 1 | 4.99 | 4.81 | 4.57 | 4.83 | 4.84 | 4.89 | 4.59 | 4.73 | 4.69 | 5.00 | 4.72 | 4.77 | 4.74 |
| Biotic bottle 2 | 4.87 | 4.93 | 4.82 | 4.58 | 4.46 | 4.90 | 4.82 | 4.80 | 4.77 | 4.76 | 4.75 | 4.84 | 4.74 |
| Biotic bottle 3 | 4.88 | 4.65 | 4.86 | 4.91 | 4.83 | 4.88 | 4.82 | 4.82 | 4.83 | 4.76 | 4.84 | 4.76 | 4.72 |
| Biotic bottle 4 | 4.91 | 4.36 | 4.86 | 4.87 | 4.93 | 4.94 | 4.88 | 4.84 | 4.71 | 4.75 | 4.64 | 4.72 | 4.85 |
| Abiotic control MEAN (n=3) | 4.91 | 4.77 | 4.90 | 4.87 | 4.93 | 4.92 | 4.88 | 4.88 | 4.84 | 4.73 | 4.81 | 4.88 | 4.83 |

<LOQ: below the limit of quantificaiton.

LOQ for volatile fatty acids and inorganic aions (mmol/L): Lactate: 0.003; Acetate: 0.001; Propionate: 0.001; Formate:0.001; Butyrate: 0.001; Pyruvate: 0.001; Chloride: 0.004; Nitrite: 0.002; Nitrate: 0.004; Sulfate: 0.001; Phosphate: 0.002.

**Table S7. Data for headspace methane analysis (µmol/L) and calculated methane production**

| | | | Biotic bottle 1 | | Biotic bottle 2 | | Biotic bottle 3 | | Biotic bottle 4 | | Abiotic control MEAN (n=3) |
|---|---|---|---|---|---|---|---|---|---|---|---|
| Day | Liquid volume (L) | Headspace volume (L) | Headspace methane (µmol/L) | Mathane production (µmol/bottle) | Headspace methane (µmol/L) | Mathane production (µmol/bottle) | Headspace methane (µmol/L) | Mathane production (µmol/bottle) | Headspace methane (µmol/L) | Mathane production (µmol/bottle) | Headspace methane (µmol/L) |
| 0 | 0.210 | 0.040 | <LOQ | 0 | <LOQ | 0 | <LOQ | 0 | <LOQ | 0 | <LOQ |
| 2 | 0.209 | 0.041 | 120 | 6 | 98 | 5 | 99 | 5 | 80 | 4 | <LOQ |
| 5 | 0.208 | 0.042 | 873 | 43 | 919 | 45 | 995 | 49 | 844 | 41 | <LOQ |
| 9 | 0.207 | 0.043 | 1017 | 51 | 1532 | 77 | 1047 | 53 | 861 | 43 | <LOQ |
| 19 | 0.201 | 0.049 | 881 | 49 | 1244 | 70 | 871 | 49 | 788 | 44 | <LOQ |
| 28 | 0.195 | 0.055 | 913 | 56 | 1245 | 77 | 838 | 52 | 824 | 51 | <LOQ |
| 39 | 0.189 | 0.061 | 1152 | 78 | 1261 | 86 | 828 | 56 | 783 | 53 | <LOQ |
| 49 | 0.182 | 0.068 | 1284 | 95 | 1022 | 76 | 791 | 59 | 755 | 56 | <LOQ |
| 79 | 0.172 | 0.078 | 1624 | 137 | 1168 | 99 | 1180 | 100 | 892 | 75 | <LOQ |
| 94 | 0.163 | 0.087 | 1788 | 166 | 1298 | 121 | 1266 | 118 | 1226 | 114 | <LOQ |

<LOQ: below the limit of quantificaiton. LOQ for headspace methane is 35 µmol/L.

**Table S8. Summary of measured growth yields of *Dehalobacter* (Dhb) in various units.**

| Growth yield on 2,3-DCA | | | Growth yield on 2-CA | | |
|---|---|---|---|---|---|
| Copies per µmol chloride released | Dhb cells per mol chloride released | g Dhb per g 2,3-DCA | Copies per µmol chloride released | Dhb cells per mol chloride released | g Dhb per g 2-CA |
| $1.2 \pm 0.1 \times 10^{8}$ | $3.0 \pm 0.3 \times 10^{13}$ | $1.7 \pm 0.2 \times 10^{-2}$ | $1.3 \pm 0.1 \times 10^{8}$ | $3.3 \pm 0.2 \times 10^{13}$ | $2.4 \pm 0.2 \times 10^{-2}$ |

Note: Yields in the unit of Dhb cells per mol chloride released can be used for comparison with previously reported values summarized in a companion work.[1]

Assumptions for unit conversion:

1. Each *Dehalobacter* cell was assumed to correspond to four 16S rRNA gene copies.

2. To maximize the utility of the results for bioremediation and compare the measured yields with theoretical yields, a computation approach as illustrated in Qiao et al. (2018),[11] which considers the cellular shape and cell volume of *Dehalobacter*, was employed to estimate the *Dehalobacter* biomass concentration in gram cell per litre, assuming a biomass for *Dehalobacter* of $9.10 \times 10^{-14}$ g per dry cell.[11]

**Table S9. Comparison of measured and predicted growth yields for *Dehalobacter* (Dhb) at different values of the energy transfer efficiency (ɛ̧).**

| Growth yield on 2,3-DCA | | | | Growth yield on 2-CA | | | |
|---|---|---|---|---|---|---|---|
| $\Delta G_r^{o'}$ | -68.63 kJ/e⁻eq | | | $\Delta G_r^{o'}$ | -64.53 kJ/e⁻eq | | |
| $f_s$ | 0.53 e⁻eq Dhb cell/ e⁻eq $H_2$ | | | $f_s$ | 0.51 e⁻eq Dhb cell/ e⁻eq $H_2$ | | |
| Measured yield (g Dhb cells / g 2,3-DCA) | Energetics model | | | Measured yield (g Dhb cells / g 2-CA) | Energetics model | | |
| | ε | Predicted yield (g Dhb cells / g 2,3-DCA) | Predicted yield (Dhb cells / mol 2,3-DCA) | | ε | Predicted yield (g Dhb cells / g 2-CA) | Predicted yield (Dhb cells / mol 2-CA) |
| $1.7 \pm 0.2\times10^{-2}$ | 60% | $4.1\times10^{-2}$ | $7.3\times10^{13}$ | $2.4 \pm 0.2\times10^{-2}$ | 60% | $4.7\times10^{-2}$ | $6.6\times10^{13}$ |
| | 56% | $3.3\times10^{-2}$ | $5.9\times10^{13}$ | | 56% | $3.8\times10^{-2}$ | $5.3\times10^{13}$ |
| | 54% | $3.0\times10^{-2}$ | $5.3\times10^{13}$ | | 54% | $3.4\times10^{-2}$ | $4.8\times10^{13}$ |
| | 52% | $2.6\times10^{-2}$ | $4.6\times10^{13}$ | | 52% | $3.0\times10^{-2}$ | $4.2\times10^{13}$ |
| | 50% | $2.3\times10^{-2}$ | $4.1\times10^{13}$ | | 50% | $2.7\times10^{-2}$ | $3.8\times10^{13}$ |
| | 48% | $2.1\times10^{-2}$ | $3.7\times10^{13}$ | | 48% | $2.4\times10^{-2}$ | $3.4\times10^{13}$ |
| | 45% | $1.7\times10^{-2}$ | $3.0\times10^{13}$ | | 45% | $1.9\times10^{-2}$ | $2.7\times10^{13}$ |
| | 43% | $1.4\times10^{-2}$ | $2.5\times10^{13}$ | | 43% | $1.7\times10^{-2}$ | $2.4\times10^{13}$ |
| | 40% | $1.1\times10^{-2}$ | $2.0\times10^{13}$ | | 40% | $1.3\times10^{-2}$ | $1.8\times10^{13}$ |

The calculations and assumptions for yield predictions based on the energetics model are detailed in Section S9. The $f_s$ values shown in the table were calculated assuming an ɛ̧ of 60%.

**Table S10. Evaluation of linear regressions for calculating the growth yields of *Dehalobacter* associated with 2,3-DCA dechlorination (days 0 to 50) and 2-CA dechlorination (days 50 to 100).**

| | Yield (slope) (copies/ µmol 2,3-DCA or 2-CA consumed) [a] | Std error [b] on Y | 95%CI[c] on Y | p-value of Y | F-statistic | $R^2$ | Sample size |
|---|---|---|---|---|---|---|---|
| **2,3-DCA dechlorination** | $1.22\times10^{8}$ | $1.14\times10^{7}$ | $2.23\times10^{7}$ | $3.11\times10^{-9}$ | 114.5 | 0.86 | 20 |
| **2-CA dechlorination** | $1.33\times10^{8}$ | $9.69\times10^{6}$ | $1.90\times10^{7}$ | $9.17\times10^{-6}$ | 189.3 | 0.97 | 8 |

[a]Yield: The growth yield associated with 2-CA dechlorination was calculated using the aniline production, leading to an original unit of copies per µmol aniline formed. As one mole of aniline formation led to one mole of 2-CA consumed, this unit is the same as copies per µmol 2-CA consumed. This unit is also equivalent to copies per µmol chloride released.

[b] Std error: Standard error

[c] 95%CI: 95% confidence interval, calculated as 1.96×Std error, assuming a normal distribution of the sample mean.

**Table S11. Results and evaluation of the Monod kinetics to determine Monod constants ($K_s$ and $\mu_{max}$).**

| | 2,3-DCA dechlorination by *Dehalobacter* | 2-CA dechlorination by *Dehalobacter* |
|---|---|---|
| Constants used for the best fit: | | |
| Y (g/g) | $1.7 \times 10^{-2}$ | $2.4 \times 10^{-2}$ |
| $k_d$ ($day^{-1}$) | 0.017 | |
| $X_0$' (g/L) | $8.3 \times 10^{-5}$ | |
| Obtained Monod parameters: | | |
| $K_s \pm$ SE (mg/L) | 45 ± 16 | 35 ± 24 |
| $\mu_{max} \pm$ SE ($day^{-1}$) | 0.18 ± 0.03 | 0.14 ± 0.06 |
| Model evaluation: | | |
| RSS | $7.71\times 10^{-4}$ | $5.08\times 10^{-5}$ |
| Reduced chi-squared | 1.1 | 1 |
| RMSE | $2.82\times 10^{-3}$ | $1.45\times 10^{-3}$ |

SE: standard error.

#### Table S12. Calculation of electron balances and distributions

| Biotic bottle 1 | | | | | | | | | | |
|---|---|---|---|---|---|---|---|---|---|---|
| Day | total methane (μmol e- ) | dechlorination I (μmol e- ) | dechlorination II (μmol e- ) | ethanol (μmol e- ) | lactate (μmol e- ) | acetate (μmol e- ) | propionate (μmol e- ) | formate (μmol e- ) | Hydrogen (μmol e- ) | sum (μmol e- ) |
| 0 | 0.0 | 0.0 | 0.0 | 8316* | 1584.7 | 388.5 | 0.3 | 14.9 | 39.6* | 10344.1 |
| 5 | 341.9 | 5.7 | 0.0 | 0* | 0.0 | 5614.8 | 1214.3 | 20.1 | N.A. | 7196.8 |
| 9 | 410.3 | 11.9 | 0.0 | 0* | 0.0 | 5603.0 | 1198.1 | 50.8 | N.A. | 7274.2 |
| 22 | 395.8 | 42.1 | 1.7 | 0* | 0.0 | 5744.4 | 1219.8 | 13.4 | N.A. | 7417.1 |
| 29 | 451.3 | 58.3 | 2.4 | 0* | 0.0 | 5859.3 | 1203.5 | 33.8 | N.A. | 7608.6 |
| 36 | 625.5 | 76.7 | 2.1 | 0* | 0.0 | 5638.3 | 1203.5 | 22.9 | N.A. | 7569.1 |
| 50 | 760.6 | 97.5 | 5.3 | 0* | 0.0 | 5783.9 | 933.8 | 21.0 | N.A. | 7602.0 |
| 79 | 1096.9 | 97.9 | 87.1 | 0* | 0.0 | 5803.7 | 511.4 | 23.4 | N.A. | 7620.4 |
| 94 | 1331.6 | 97.9 | 104.4 | 0* | 0.0 | 5919.1 | 83.0 | 12.0 | N.A. | 7548.1 |
| | | | | | | | eeq recovery | | | 72.97% |
| | | | | From days 50 to 94, methane increase / propionate decrease is | | | | | | 67.1% |
| Biotic bottle 2 | | | | | | | | | | |
| Day | total methane (μmol e- ) | dechlorination I (μmol e- ) | dechlorination II (μmol e- ) | ethanol* (μmol e- ) | lactate (μmol e- ) | acetate (μmol e- ) | propionate (μmol e- ) | formate (μmol e- ) | Hydrogen (μmol e- ) | sum (μmol e- ) |
| 0 | 0.0 | 0.0 | 0.0 | 8316* | 1537.0 | 354.0 | 0.3 | 10.6 | 39.6* | 10257.4 |
| 5 | 359.8 | 5.8 | 0.0 | 0* | 0.0 | 7603.1 | 1225.2 | 16.3 | N.A. | 9210.1 |
| 9 | 617.8 | 10.4 | 0.0 | 0* | 0.0 | 7648.4 | 1219.8 | 39.0 | N.A. | 9535.3 |
| 22 | 558.4 | 29.8 | 0.0 | 0* | 0.0 | 6993.5 | 1117.0 | 12.0 | N.A. | 8710.7 |
| 29 | 615.7 | 40.2 | 0.0 | 0* | 0.0 | 7780.7 | 1214.3 | 20.1 | N.A. | 9671.0 |
| 36 | 684.3 | 55.2 | 0.0 | 0* | 0.0 | 7812.8 | 1019.9 | 40.4 | N.A. | 9612.6 |
| 50 | 605.4 | 85.2 | 1.7 | 0* | 0.0 | 7725.8 | 960.7 | 26.7 | N.A. | 9405.4 |
| 79 | 788.4 | 99.5 | 10.5 | 0* | 0.0 | 7352.2 | 751.4 | 16.3 | N.A. | 9018.3 |
| 94 | 966.4 | 99.5 | 27.0 | 0* | 0.0 | 7575.9 | 506.1 | 15.8 | N.A. | 9190.7 |
| | | | | | | | eeq recovery | | | 89.60% |
| | | | | From days 50 to 94, methane increase / propionate decrease is | | | | | | 79.4% |
| Biotic bottle 3 | | | | | | | | | | |
| Day | total methane (μmol e- ) | dechlorination I (μmol e- ) | dechlorination II (μmol e- ) | ethanol* (μmol e- ) | lactate (μmol e- ) | acetate (μmol e- ) | propionate (μmol e- ) | formate (μmol e- ) | Hydrogen (μmol e- ) | sum (μmol e- ) |
| 0 | 0.0 | 0.0 | 0.0 | 8316* | 1587.7 | 365.5 | 0.3 | 10.1 | 39.6* | 10319.2 |
| 5 | 389.5 | 7.6 | 0.0 | 0* | 0.0 | 7671.1 | 1350.0 | 8.2 | N.A. | 9426.3 |
| 9 | 422.5 | 13.4 | 0.0 | 0* | 0.0 | 9644.0 | 1225.2 | 50.4 | N.A. | 11355.4 |
| 22 | 391.1 | 40.4 | 2.0 | 0* | 0.0 | 9584.9 | 1225.2 | 35.2 | N.A. | 11278.8 |
| 29 | 414.5 | 53.5 | 2.0 | 0* | 0.0 | 9779.6 | 1225.2 | 37.6 | N.A. | 11512.4 |
| 36 | 449.5 | 69.2 | 3.0 | 0* | 0.0 | 9905.9 | 1079.2 | 22.0 | N.A. | 11528.7 |
| 50 | 468.7 | 98.8 | 10.4 | 0* | 0.0 | 9779.6 | 1052.2 | 22.4 | N.A. | 11432.1 |
| 79 | 796.5 | 99.4 | 57.3 | 0* | 0.0 | 9143.8 | 917.7 | 20.5 | N.A. | 11035.1 |
| 94 | 943.1 | 99.4 | 82.9 | 0* | 0.0 | 8934.7 | 864.0 | 19.1 | N.A. | 10943.3 |
| | | | | | | | eeq recovery | | | 106.05% |
| | | | | Only a small reduction in propionate was observed | | | | | | |
| Biotic bottle 4 | | | | | | | | | | |
| Day | total methane (μmol e- ) | dechlorination I (μmol e- ) | dechlorination II (μmol e- ) | ethanol (μmol e- ) | lactate (μmol e- ) | acetate (μmol e- ) | propionate (μmol e- ) | formate (μmol e- ) | Hydrogen (μmol e- ) | sum (μmol e- ) |
| 0 | 0.0 | 0.0 | 0.0 | 8316* | 1761.0 | 354.0 | 0.3 | 7.8 | 39.6* | 10478.6 |
| 5 | 330.3 | 5.4 | 0.0 | 0* | 0.0 | 855.6 | 1073.8 | 25.8 | N.A. | 2290.9 |
| 9 | 347.2 | 12.3 | 0.0 | 0* | 0.0 | 1045.6 | 1507.9 | 29.1 | N.A. | 2942.0 |
| 22 | 353.6 | 52.3 | 0.0 | 0* | 0.0 | 1455.6 | 1507.9 | 18.7 | N.A. | 3388.0 |
| 29 | 407.7 | 68.5 | 0.9 | 0* | 0.0 | 1673.3 | 1431.6 | 31.0 | N.A. | 3612.8 |
| 36 | 425.2 | 85.9 | 1.7 | 0* | 0.0 | 1794.0 | 1279.4 | 35.7 | N.A. | 3621.9 |
| 50 | 447.1 | 100.1 | 3.2 | 0* | 0.0 | 1956.1 | 1214.3 | 14.9 | N.A. | 3735.7 |
| 79 | 602.2 | 100.1 | 101.7 | 0* | 0.0 | 3288.2 | 890.8 | 23.4 | N.A. | 5006.4 |
| 94 | 912.7 | 100.1 | 105.1 | 0* | 0.0 | 5194.5 | 687.3 | 20.5 | N.A. | 7020.3 |
| | | | | | | | eeq recovery | | | 67.00% |
| | | | | From days 50 to 94, methane increase / propionate decrease is | | | | | | 88.3% |

All values not directly measured are marked with an asterisk (*). Electrons in ethanol at time zero were calculated based on the theoretical amount added to the system. Ethanol was assumed to be fully depleted by day 5. Initial headspace hydrogen was estimated from the hydrogen concentration in the glovebox atmosphere. N.A.: not measured
